# The *Shigella* E3 ubiquitin ligase IpaH7.8 reprograms host kinase signaling to suppress NOX2-dependent oxidative burst responses in human monocytes

**DOI:** 10.64898/2026.04.28.721327

**Authors:** Theresa Karagöz, Ivan Ngueya Yango, Stefanie Norkowski, Yannick Teschke, Britta Körner, Joyleen Fernandes, Yvonne Boergeling, Petra Dersch, Christian Rüter

**Author notes:** Corresponding author: Christian Rüter, Institute for Infectiology, Center for Molecular Biology of Inflammation (ZMBE), Von Esmarch Str. 56, 48149 Münster, Germany.

## Abstract

Bacterial effector proteins manipulate host signalling cascades, including immune responses, to facilitate infection. While most effectors of Gram-negative bacteria rely on a secretion system for intracellular delivery, some possess intrinsic cell-penetrating capabilities. Here, we characterize the *Shigella flexneri* LPX effector IpaH7.8, which combines autonomous cell entry with enzymatic modulation of immunomodulatory host signaling pathways through distinct structural domains. We show that recombinant IpaH7.8 (rIpaH7.8) enters human cells independent of *Shigella*’s type III secretion system (T3SS) via lipid raft-mediated endocytosis and escapes the endosome through a conserved N-terminal domain composed of two α-helices. In the cytosol, the C-terminal E3 ubiquitin ligase domain of the cell-penetrating effector protein targets the pore-forming protein gasdermin D (GSDMD), suppressing inflammasome-induced IL-1β release. Beyond inflammasome inhibition, integrated transcriptomic and kinome profiling in primary human monocytes revealed that IpaH7.8 induces a coordinated reprogramming of host signaling networks. Cluster-resolved gene expression analysis demonstrated selective suppression of immune effector pathways alongside induction of regulatory programs and interference with vesicular trafficking. These transcriptional changes converged with kinase activity remodeling, characterized by attenuation of PKC- and PKA-dependent signaling pathways. Notably, both datasets identified the NOX2 complex as a central target of IpaH7.8 activity. The NOX2 subunit NCF1 was downregulated at the transcriptional level and showed reduced phosphorylation at regulatory sites, indicating impaired activation. Consistently, IpaH7.8 significantly reduced reactive oxygen species production in primary human monocytes, demonstrating functional suppression of oxidative burst responses.

Together, our findings reveal that IpaH7.8 acts as a multi-layered regulator of host immunity that integrates ubiquitination and kinase signaling to suppress both inflammatory and antimicrobial responses. By converging on the NOX2 axis, this effector uncovers a central vulnerability in host defense and highlights bacterial effector proteins as modulators of complex signaling networks with potential therapeutic relevance.

**Author Summary:** Bacterial pathogens like *Shigella flexneri* manipulate host immune responses to survive and spread within human cells. The *Shigella* effector protein IpaH7.8 is known to block inflammatory cell death by targeting gasdermin D. Here, we show that IpaH7.8 can enter human cells without a bacterial secretion system. It uses a specialized protein domain to cross the membrane and reach the cytoplasm. Once inside, IpaH7.8 alters host cell signaling by both attaching ubiquitin to immune proteins and reprogramming phosphorylation pathways. This dual function allows *Shigella* to suppress inflammation and promotes its escape from immune defenses. Our findings reveal how IpaH7.8 combines cell entry, immune evasion, and cytoskeletal control in a single protein, and highlight its potential as a tool to modulate inflammation in disease contexts.

## Introduction

*Shigella flexneri* is a Gram-negative, facultative intracellular pathogen and a major cause of bacillary dysentery (shigellosis), a severe intestinal infection characterized by inflammation, abdominal pain, diarrhea, and fever. Shigellosis accounts for over 270 million infections worldwide annually, disproportionately affecting children in low-income regions with limited access to clean water and sanitation [1,2]. The primary mode of transmission is through the ingestion of contaminated food or water, resulting in the colonization and invasion of the intestinal epithelium. After ingestion, *S. flexneri* translocates through specialized microfold (M) cells in the intestinal epithelium, where it is phagocytosed by macrophages and dendritic cells. Upon uptake, the bacterium rapidly escapes from the bacteria-containing vacuole (BCV) into the cytosol, where it replicates and directly interacts with the host immune system [1,2]. To evade host immune responses and establish an intracellular replicative niche, *S. flexneri* utilizes a type III secretion system (T3SS) to inject bacterial effector proteins into host cells, modulating immune signaling and cell death pathways [3–6]. Among these effectors, the IpaH proteins, a unique class of E3 ubiquitin ligases classified as LPX effectors (harboring a **L**eu-**P**ro-**X** motif), play a crucial role in subverting the host’s innate immune defenses by targeting key regulators of immune signaling, pyroptosis, and inflammasome activation [7]. Pyroptosis is a pro-inflammatory form of regulated cell death that serves as a crucial immune defense mechanism against intracellular bacterial infections. This process is primarily mediated by the gasdermin family of pore-forming proteins, particularly gasdermin D (GSDMD), which, upon activation, oligomerizes and forms membrane pores, leading to host cell lysis and the release of inflammatory cytokines such as interleukin-1 beta (IL-1β) and interleukin-18 (IL-18) [4,8]. The *Shigella* effector IpaH7.8 ubiquitinates and targets GSDMD for proteasomal degradation, thereby inhibiting pyroptosis and enabling bacterial survival in the cytosol [9]. Notably, IpaH7.8 displays species-specific activity, efficiently targeting human GSDMD but not murine GSDMD due to structural differences. This specificity may contribute to the restricted host range of *S. flexneri*, which primarily infects humans and non-human primates but not mice [9]. Additionally, IpaH7.8 targets gasdermin B (GSDMB), another pore-forming cytolysin, preventing granzyme A-mediated activation in epithelial cells and thereby protecting *Shigella* from the bactericidal effects of natural killer cells [10,11]. Beyond gasdermin regulation, IpaH7.8 has also been implicated in inflammasome modulation through its interaction with glomulin (GLMN), a negative regulator of inflammasome activation. The degradation of GLMN by IpaH7.8 leads to the activation of NLRP3 and NLRC4 inflammasomes, promoting caspase-1-dependent pyroptosis in murine macrophages [3,4]. Additionally, IpaH7.8 triggers proteasome-mediated degradation of the N-terminal domain of murine NLRP1B, releasing a C-terminal fragment that is a potent activator of caspase-1 [12]. However, both NLRP1B and GLMN have thus far been identified only as targets of IpaH7.8 in murine cells, and their relevance in human cells remains unclear. In addition to its role in immune evasion, the *Shigella* effector protein IpaH7.8 has recently been shown to also regulate Rab13, promoting its retention on the membrane remnants of the *Shigella*-containing vacuole (SCV) [13]. This process facilitates efficient uncoating of SCV membranes, which is likely essential for IpaH7.8-dependent bacterial escape and successful intercellular spread [2]. The interaction between IpaH7.8 and Rab13 appears to be unique among the IpaH family members, with no functional redundancy observed for other bacterial effectors [13]. Despite extensive research, the precise molecular function and role of IpaH7.8 in different host cell types, as well as its broader implications for *Shigella* pathogenesis, warrants further investigation. Moreover, the precise mechanisms underlying their translocation and functional impact on host cell signaling remain unresolved. Our previous studies demonstrated that LPX effectors can penetrate host cells independently of the bacterial T3SS, a property shared by multiple family members, including IpaH proteins from *Shigella flexneri*, SspH proteins from *Salmonella*, and YopM from *Yersinia* [14–16]. With the exception of IpaH9.8 [15], the mechanisms underlying autonomous cell entry of other IpaH proteins, including IpaH7.8, remain poorly understood. Here, we focus on IpaH7.8, an LPX effector composed of N-terminal α-helices followed by central leucine-rich repeat (LRR) domains containing LPX consensus motifs (Fig 1A). These LRR domains form a characteristic curved, horseshoe-like structure and are primarily involved in protein-protein interactions [7,17]. In contrast to YopM, which lacks a defined enzymatic domain, IpaH7.8 harbors a conserved C-terminal α-helical domain that functions as a novel E3 ubiquitin ligase (NEL) [18]. The N-terminus of LPX effectors contains a flexible region followed by two conserved α-helices (Fig 1B and 1D), which structurally resemble cell-penetrating peptides (CPPs). Notably, unlike classical cationic CPPs such as Tat (Trans-Activator of Transcription, human immunodeficiency virus 1) [20], these protein transduction domains (PTDs) are negatively charged at physiological pH and therefore classify as anionic CPPs (Fig 1B and 1C). A well-characterized example of this class is the sweet-arrow peptide Sap(E), which enters cells via aggregation and endocytosis and exhibits pH-dependent membrane interaction [21,22]. Similarly, LPX PTDs display a pH-dependent shift toward increased positive charge under mildly acidic conditions, suggesting that they function as dynamic, pH-sensitive modules facilitating cellular entry (Fig 1C). However, particularly for IpaH7.8, the molecular basis and functional consequences of this N-terminal domain mediating autonomous entry remain largely unresolved. To address this, the present study focuses on the *Shigella* effector IpaH7.8. We systematically characterize its T3SS-independent translocation into different host cell types using deletion constructs and recombinant protein approaches. In addition, we identify and functionally validate N-terminal protein transduction domains that mediate cellular uptake. Beyond entry, we investigate whether IpaH7.8 retains its enzymatic activity as an E3 ubiquitin ligase following cell penetration and assess its impact on host signaling pathways. By linking the cell-penetrating capacity of IpaH7.8 to its ability to reprogram host immune signaling, this study provides new mechanistic insight into the role of LPX effectors during Shigella infection.

**Figure 1:**
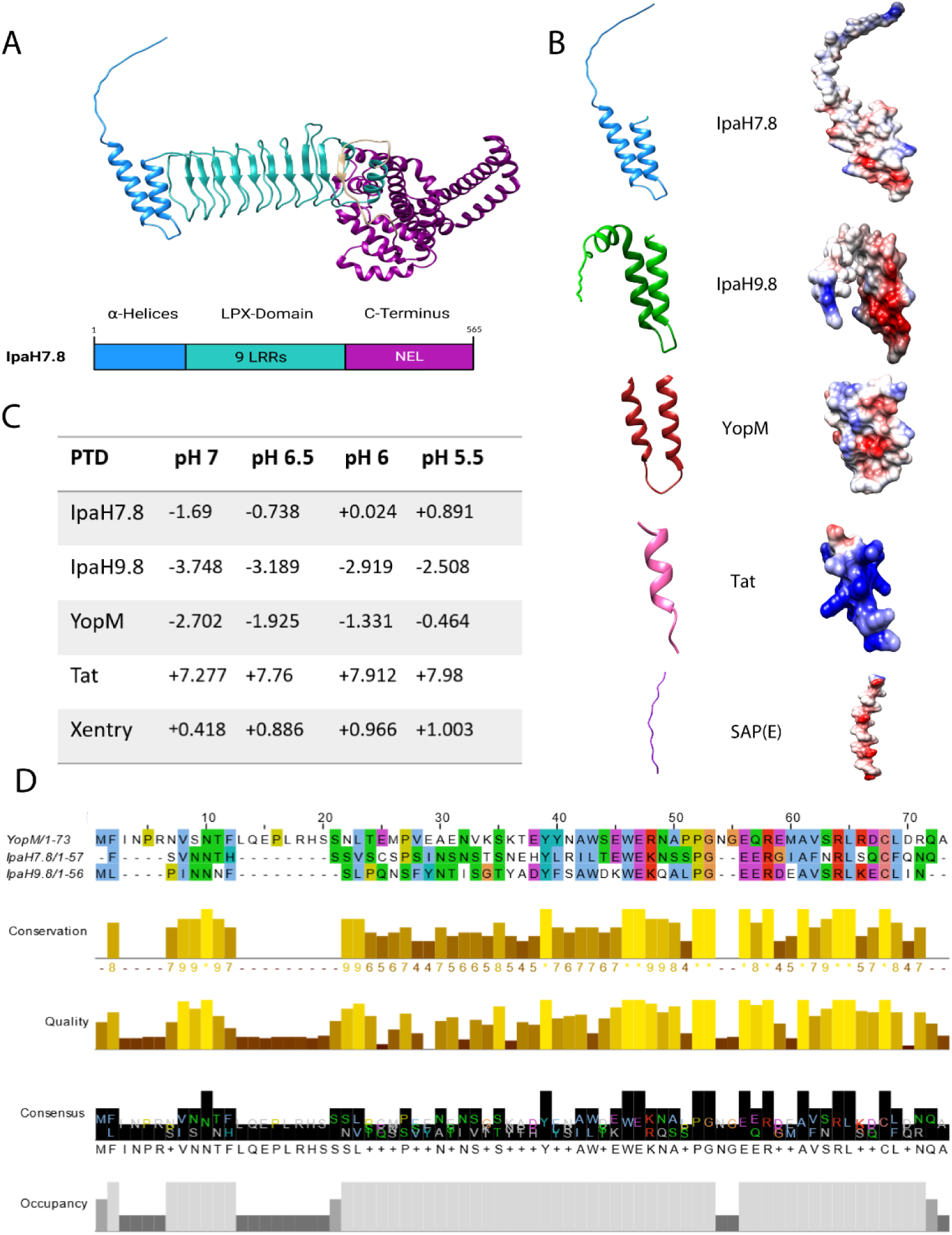
Structural visualization of IpaH7.8 and comparison to bacterial LPX effector proteins YopM from *Yersinia enterocolitica* and IpaH9.8 from *Shigella flexneri*. (A) AlphaFold-predicted ribbon model of full-length IpaH7.8 (*Shigella flexneri;* UniProt: P18014) visualized in UCSF Chimera. The C-terminal novel E3 ubiquitin ligase (NEL) domain structure corresponds to a high-confidence prediction from the AlphaFold Protein Structure Database (P18014). The schematic representation below shows the domain architecture of IpaH7.8 (strain M90T), with N-terminal domains in blue, LPX domains in green, and the E3 ubiquitin ligase domain in purple. (B) Surface representations of the N-terminal α-helical domains of YopM (AA_1-73_), IpaH9.8 (AA_1-56_), IpaH7.8 (AA_1-57_), Tat (MGYGRKKRRQRRRG), and SAP(E) (CGGWVELPPPVELPPPVELPPP). Charged amino acid residues are colored according to electrostatic potential (coulombic surface coloring) in blue (pos.) and red (neg.), respectively. (C) Calculated net charge of indicated protein transduction domains (PTDs) at different pH. (D) Alignment of N-terminal domains of IpaH7.8, IpaH9.8, and YopM. The alignment was obtained with MUSCLE with default settings and visualized using JalView.

## Results

### rIpaH7.8 is a cell-penetrating effector protein with a functional N-terminal protein transduction domain for cytosolic transport

To analyze the T3SS-independent uptake of the recombinant LPX effector protein, microscopy studies and a flow cytometry-based assay were performed. The latter used fluorescent dye-conjugated recombinant IpaH7.8 (rIpaH7.8), as previously established [14,15]. Upon incubation of HeLa cells with FITC-labeled rIpaH7.8, confocal microscopy visualized rIpaH7.8 within the cytosol of HeLa cells and revealed a predominant perinuclear accumulation pattern, suggesting endocytic uptake followed by intracellular redistribution (Fig 2A and S1B Fig). To specifically quantify intracellular fluorescence, trypan blue was employed to quench extracellular- and surface-associated signals [16]. In this regard, a pronounced dose- and time-dependent increase in intracellular fluorescence was detected by fluorescence-based flow cytometry, comparable to that observed for the cell-penetrating effectors (CPEs) rYopM and rIpaH9.8 (Fig 2C, S1A Fig and S1C-D) [14,17]. Similar results were obtained in HEK293T epithelial cells and THP-1-derived macrophages, confirming that this T3SS-independent internalization mechanism is not restricted to a specific cell type (S1E-J Fig). To define the structural determinants responsible for cellular translocation, we next examined uptake of rIpaH7.8 deletion constructs (Fig 2B). The C-terminal truncation mutant (ΔC), which lacks E3 ubiquitin ligase (NEL) activity, was efficiently internalized, whereas the N-terminal deletion variant (ΔN) displayed markedly reduced uptake (Fig 2A and 2C, S1H Fig and S1J). These results identify the N-terminal α-helical region of IpaH7.8 as a functional protein transduction domain (PTD), as demonstrated for other CPEs of the LPX family [14,18,19]. To test whether this region alone can transport heterologous cargo, we generated a fusion construct of the N-terminal 2α-helices of IpaH7.8 linked to GFP (S2A Fig). This 2αH-IpaH7.8-GFP construct was efficiently taken up by host cells, comparable to the positive control 2αH-YopM-GFP [20], whereas recombinant GFP alone showed only background fluorescence (S2B-C Fig). To elucidate the mechanism of T3SS-independent translocation, we examined the energy dependence and endocytic requirements of rIpaH7.8 uptake. Internalization was markedly reduced at 4°C, indicating that the process depends on active endocytosis rather than passive diffusion (S2D Fig). Pharmacological inhibition revealed that Cytochalasin D and Amiloride, which interfere with actin polymerization and macropinocytosis, significantly impaired uptake, whereas inhibitors of clathrin-mediated or dynamin-dependent endocytosis (Dynasore) had only moderate effects (Fig 2E). Likewise, disruption of lipid rafts by Methyl-β-cyclodextrin (MβCD) diminished internalization, implicating lipid raft-dependent macropinocytosis as the major entry pathway (Fig 2E). Co-localization analyses with fluorescent endocytic tracers supported these results, showing preferential association of rIpaH7.8 with the cholera toxin B subunit (CTB), a marker of raft-dependent uptake [21], rather than with transferrin, a marker of clathrin-dependent endocytosis (S2E Fig) [22]. Following entry, rIpaH7.8 sequentially localized to Rab5-positive early endosomes, Rab7-positive late endosomes, and CD63-positive lysosomes, suggesting endosomal trafficking through the canonical endolysosomal pathway (Fig 2D). However, endosomal escape assays using the pH-sensitive dye naphthofluorescein (NF, S2F Fig) [23], demonstrated that a substantial fraction of rIpaH7.8 reached the cytosol, as indicated by fluorescence activation in neutral intracellular environments (Fig 2F and S2G Fig). Subcellular fractionation further confirmed the presence of rIpaH7.8 in the cytoplasmic fraction, indicating successful release from endosomal compartments like other CPEs of the LPX subtype (Fig 2G), [14,17,20,24]. However, rIpaH7.8, similar to rIpaH9.8, was present in the membrane fraction in comparable amounts (Fig 2G), suggesting that a portion of the proteins remained membrane-associated or is trapped within endosomal compartments. Together, these results demonstrate that rIpaH7.8 is a CPE that can enter cells via macropinocytosis and lipid-raft-dependent pathways, escapes from endosomes to reach the cytosol, and uses its N-terminal α-helical domain for uptake and cargo delivery.

**Figure 2.**
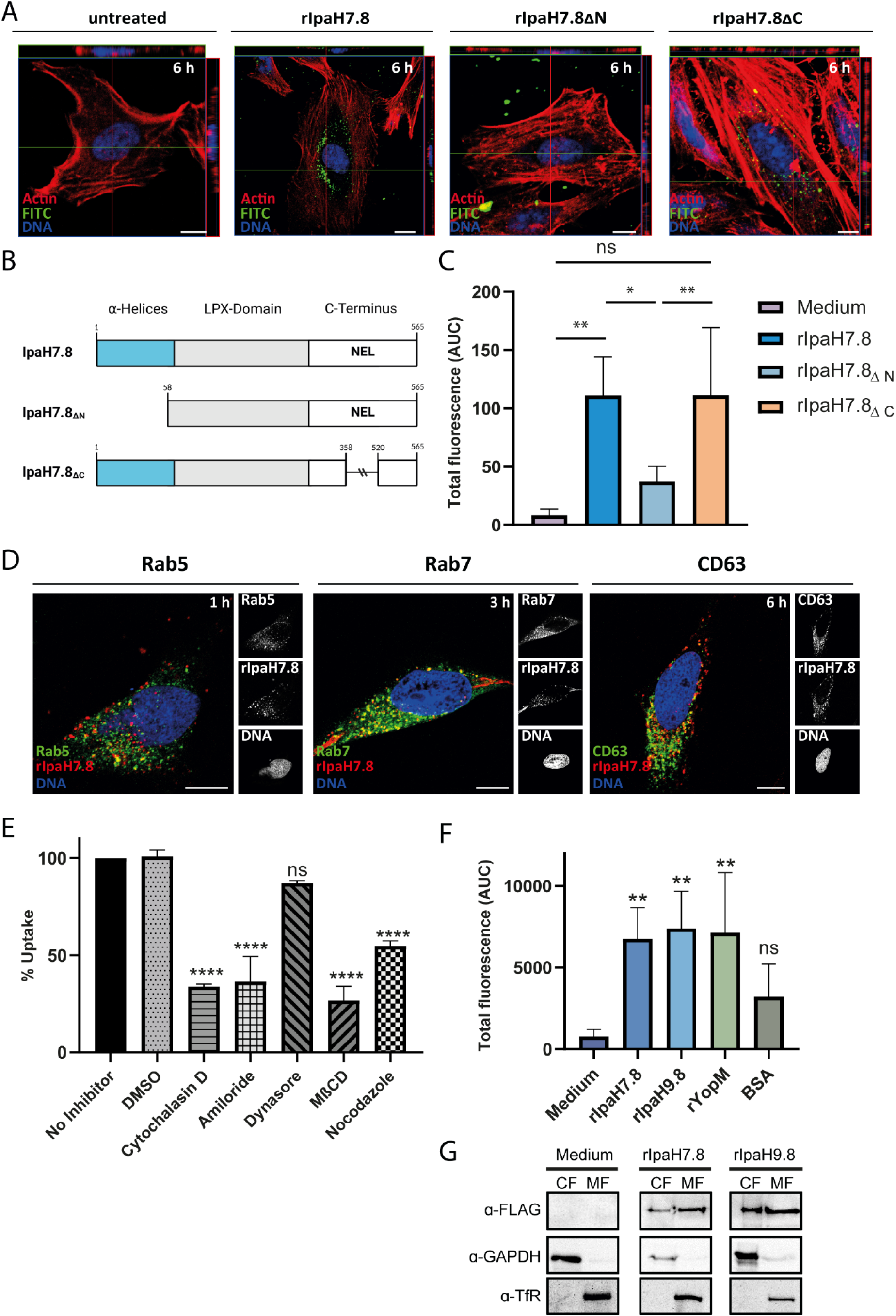
T3SS-independent uptake and intracellular trafficking of recombinant IpaH7.8. (A) Representative fluorescence microscopy images of HeLa cells incubated for 6 h with 0.4 µM FITC-labeled rIpaH7.8 or deletion variants (ΔN, ΔC). Actin (red), FITC-labeled protein (green), nuclei (blue). Scale bar, 10 µm. (B) Schematic representation of IpaH7.8 domain organization and deletion constructs. NEL, novel E3 ubiquitin ligase domain. (C) Quantification of intracellular fluorescence by flow cytometry after incubation with indicated proteins for up to 6 h. Medium-only control included. Data are shown as mean ± SD (n = 3); AUC values are indicated. Statistical analysis by one-way ANOVA with Tukey’s post hoc test. ns: non-significant, *p < 0.05, **p < 0.01. (D) Intracellular trafficking of Cy3-labeled rIpaH7.8 visualized by co-localization with Rab5-, Rab7-, and CD63-positive compartments at indicated time points. Scale bar, 10 µm. All images consist of a single optical section from a z-series with a pinhole diameter of 1 airy unit. (E) Effect of endocytosis inhibitors on rIpaH7.8 uptake. HeLa cells were pretreated with inhibitors targeting macropinocytosis, actin polymerization, or lipid raft integrity prior to incubation with Alexa Fluor 488-labeled rIpaH7.8. Intracellular fluorescence was quantified by flow cytometry (mean ± SD, n = 3; one-way ANOVA with Tukey’s post hoc test, ns non-significant, *p < 0.05, ****p < 0.0001). (F) Detection of cytosolic localization using pH-sensitive naphthofluorescein-labeled rIpaH7.8 and control proteins. Intracellular fluorescence and AUC values were determined by flow cytometry (mean ± SD, n = 3; one-way ANOVA with Tukey’s post hoc test, ns non-significant, *p < 0.05 vs. BSA). (G) Immunoblot analysis of cytoplasmic (CF) and membrane (MF) fractions confirming cytosolic localization of FLAG-tagged rIpaH7.8. GAPDH and transferrin receptor (TF-R) were used as fractionation controls.

### The C-terminal E3 ubiquitin ligase (NEL) domain of rIpaH7.8 targets GSDMD to attenuate IL-1β release

Like other LPX effector proteins, rIpaH7.8 harbors a conserved C-terminal novel E3 ubiquitin ligase (NEL) domain that mediates substrate polyubiquitination [25,26], by catalyzing the attachment of ubiquitin to a target protein’s lysine residue via an isopeptide bond with ubiquitin’s C-terminal glycine (Fig 3A; [27]). To confirm that this domain is enzymatically active in recombinant rIpaH7.8, we performed *in vitro* ubiquitination assays using purified proteins. The full-length rIpaH7.8 and its N-terminal deletion variant (ΔN) catalyzed the formation of polyubiquitin chains, whereas the catalytic C-terminal deletion mutant (ΔC) lacked this activity (Fig 3B), showing that the NEL domain of rIpaH7.8 is functional. Pull-down assays using THP-1 macrophage lysates further revealed that both GLMN, a known regulatory component of SCF-type E3 complexes [3], and Gasdermin D (GSDMD), a key executor of pyroptosis [9], physically interact with rIpaH7.8 independent of its catalytic activity (Fig 3C). FLAG-tagged recombinant GFP was used as a negative control and didn’t show this interaction. To determine whether cytosolic rIpaH7.8 retains its ubiquitin ligase activity after T3SS-independent entry, we incubated THP-1 cells with rIpaH7.8 or its catalytically inactive ΔC variant and analyzed ubiquitinated targets by tandem ubiquitin-binding entity (TUBE) pull-down [14]. TUBEs covalently linked to agarose beads function as high-affinity traps for polyubiquitinated proteins, enabling their efficient enrichment from cell lysates for subsequent analysis [7,28]. To validate and compare the extent of target protein polyubiquitination, the pull-down was followed by treatment with the broad-specificity deubiquitinating enzyme USP2. The increased signal intensity of the USP2-deubiquitinated protein of interest, as detected by Western blotting, is consistent with prior enrichment of its polyubiquitinated forms [7,28]. The corresponding Western blot analysis revealed pronounced polyubiquitination of GSDMD, particularly its N-terminal pore-forming unit/fragment (PFU), in cells treated with catalytically active rIpaH7.8, whereas GLMN was unaffected (Fig 3D). This finding indicated that rIpaH7.8 directly modified GSDMD after cytosolic translocation. Functionally, this correlated with a significant reduction in IL-1β secretion following inflammasome activation by nigericin (Fig 3E), indicating that cytosolic rIpaH7.8 downregulates cytokine release through GSDMD ubiquitination rather than its interference with GLMN. In this regard, analysis of ASC::GFP reporter cells revealed that speck formation, characteristic of inflammasome assembly [29], was not affected by rIpaH7.8 treatment (Fig 3F and S3A Fig). We confirmed these effects in primary human monocytes and monocyte-derived macrophages (hMDMs) to more closely approximate the *in vivo* situation and the human host environment. Flow cytometry and confocal microscopy confirmed efficient uptake and intracellular localization of rIpaH7.8 within 3 h (Fig 3G and 3H; S3B Fig and S3C). Consistent with results from THP-1 cells, catalytically active rIpaH7.8, but not its ΔC mutant, significantly reduced IL-1β release upon LPS and Nigericin stimulation (Fig 3I). These data demonstrate that the enzymatic NEL domain of rIpaH7.8 remains functional after T3SS-independent entry, mediating cytosolic GSDMD polyubiquitination to inhibit its pore-forming function and thereby dampen inflammasome-mediated cytokine release.

**Figure 3.**
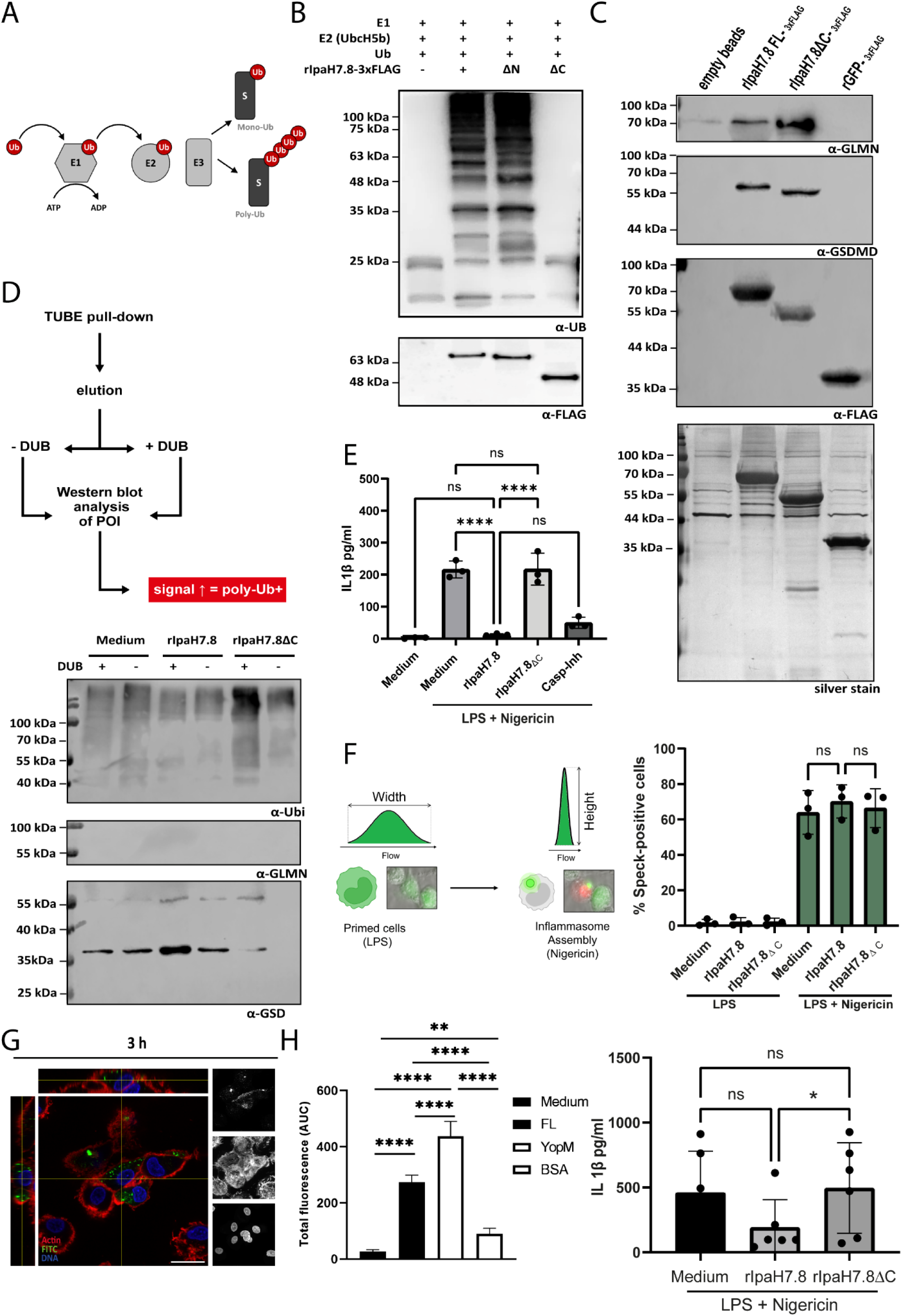
Cytosolic activity of rIpaH7.8 mediates GSDMD ubiquitination and suppresses IL-1β release. (A) Schematic overview of the ubiquitination cascade mediated by E1, E2, and E3 enzymes. (B) *In vitro* ubiquitination assay using rIpaH7.8 or deletion constructs (ΔN, ΔC), analyzed by immunoblotting with anti-ubiquitin and anti-FLAG antibodies. (C) Anti-FLAG pulldown from THP-1-derived macrophage lysates using anti-FLAG M2 magnetic beads pre-loaded with recombinant rIpaH7.8 or rIpaH7.8ΔC. rGFP-3xFLAG and unloaded beads served as controls. Note: The silver-stained loading control (rIpaHΔC) migrates at the same position in the gel as GSDMD, which may cause GSDMD to appear slightly smaller in the Western blot than its actual size. (D) TUBE pull-down from THP-1 macrophages treated with rIpaH7.8 (1.6 µM) and LPS (1 µg/mL) ± MG132, followed by Nigericin stimulation. Eluted proteins ± deubiquitinase (DUB) treatment were analyzed by immunoblotting for ubiquitin, GLMN, and GSDMD (POI, protein of interest). (E) IL-1β release from THP-1 cells primed with LPS and activated with Nigericin following pretreatment with rIpaH7.8 or rIpaH7.8ΔC. Supernatants were analyzed by ELISA. Data represents duplicates from three independent experiments; one-way ANOVA with Tukey’s post hoc test (ns: non-significant, *p < 0.05 vs. rIpaH7.8ΔC). (F) ASC speck formation in ASC::GFP-expressing THP-1 cells following LPS priming and Nigericin stimulation ± rIpaH7.8 or rIpaH7.8ΔC pretreatment. Speck-positive cells were quantified by flow cytometry using pulse-width analysis. (G) Confocal microscopy of primary human monocyte-derived macrophages incubated with FITC-labeled rIpaH7.8 (1.6 µM). Actin (red), rIpaH7.8 (green), nuclei (blue). Scale bar, 20 µm. Orthogonal projections of Merged channels (left) and maximum projection (right) show single z-sections (1 airy unit). (H) Uptake of FITC-labeled rIpaH7.8, rYopM, or BSA by primary human monocytes quantified by flow cytometry (n = 7 donors). AUC values are shown; one-way ANOVA with Tukey’s post hoc test (ns, non significant, **p < 0.01, ***p < 0.001, ****p < 0.0001). (I) IL-1β release from primary human monocytes stimulated with LPS and Nigericin ± rIpaH7.8 or rIpaH7.8ΔC pretreatment. ELISA analysis; n = 6; one-way ANOVA with Tukey’s post hoc test (ns, non significant, *p < 0.05 vs. rIpaH7.8ΔC).

### Differential gene expression analysis defines the transcriptional impact of rIpaH7.8 on primary human monocytes in four major gene expression patterns

Considering the variety of previously described pathways and processes influenced by IpaH7.8, such as inflammasome rewiring, cytokine release, but also vacuolar escape and cell-to-cell spread [2,3,9,12,13], an unbiased transcriptomic profiling was performed to evaluate the overall impact of IpaH7.8 on host gene expression in primary human monocytes. The cells were treated with rIpaH7.8 or the ΔC-variant for 6 h to ensure sufficient uptake and transcriptional changes. Simultaneously, LPS was added to the cells to induce an infection-relevant context. After incubation, total RNA was isolated and analyzed via polyA-enrichment-based bulk RNA-seq. This experimental setup enabled us to specifically attribute observed transcriptional changes to the enzymatic activity of IpaH7.8 in an immune-primed environment. Gene expression distributions were consistent across replicates, and treatment-specific transcriptional responses were evident from differential expressed gene (DEG) counts (S4A Fig). RNA-seq analysis revealed that LPS alone induced a strong transcriptional response, with > 3,300 DEGs. A similar global pattern was observed for cells treated with LPS in combination with rIpaH7.8 or its catalytic mutant rIpaH7.8ΔC (Fig 4A). In contrast, direct comparisons between LPS-only and LPS + protein treatments showed markedly fewer DEGs, indicating that addition of recombinant proteins modulates but does not override the dominant LPS response (Fig 4A). Comparison of rIpaH7.8 versus rIpaH7.8ΔC under LPS stimulated conditions identified only ∼250 DEGs, suggesting that the catalytic activity of IpaH7.8 contributes to subtle but distinct transcriptional changes (Fig 4A). A Venn diagram analysis confirmed that the majority of transcriptional changes were shared across all LPS-containing treatments, while a smaller subset of genes was specifically regulated by rIpaH7.8 or rIpaH7.8ΔC under LPS-stimulated conditions (Fig 4B). Profiles of rIpaH7.8-induced transcriptional changes are further illustrated in volcano plots (Fig 4C-E), where significantly up- and downregulated genes are highlighted in red and blue, respectively. The comparison of LPS-stimulated and rIpaH7.8-treated versus untreated cells showed a large number of significant transcriptional changes; however, this pattern largely mirrored the strong LPS-driven signature observed in the volcano plots of LPS-only treated versus untreated control cells (S4B Fig). Accordingly, the most prominently regulated genes in this comparison, including *IL1B, SLAMF7,* and *STAB*, are typical LPS-responsive markers (Fig 4C). When comparing LPS + rIpaH7.8-treated to LPS-only treated cells, the transcriptional response was less pronounced. Nonetheless, significant log₂ fold-changes were observed, ranging from −6 to +4.7 for several genes such as *RUFY4, SPP1,* and *CD24* (Fig 4D). Finally, the direct comparison between LPS-stimulated cells treated with either rIpaH7.8 or its catalytic mutant rIpaH7.8ΔC revealed a subset of DEGs attributable specifically to the enzymatic activity of the effector (Fig 4E). To detect transcriptional changes solely attributed to rIpaH7.8, we further intersected pairwise comparisons and classified differentially expressed genes into two categories: (a) NEL-dependent effects, defined as genes uniquely regulated in cells treated with catalytically active rIpaH7.8; and (b) NEL-independent effects, defined as genes similarly regulated in both rIpaH7.8- and rIpaH7.8ΔC-treated cells (S4C Fig). To account for variability across pairwise comparisons and to identify consistent catalytic effects, the datasets from all comparisons (Fig 4C-E) were integrated in an additional Venn analysis (S4D-F Fig). This approach identified 14 DEGs consistently affected across all conditions, representing the most robust rIpaH7.8-dependent transcriptional signatures (S4D Fig). To determine whether the genes identified in these intersections were not only significantly but also concordantly regulated, we further analyzed the directionality of regulation by separating up- and downregulated genes (S4E Fig and S4F). Based on these intersectional results, only genes significantly altered in the rIpaH7.8 vs. rIpaH7.8ΔC comparison under LPS stimulation were retained for downstream interpretation. These “overlapping DEGs” are displayed in a heatmap (Fig 4F; Z-scored FPKM values with Euclidean clustering, annotated version in S5 Fig). The most consistently affected genes across all comparisons are additionally listed in Supplemental Table 1. Unsupervised clustering of row-scaled gene expression values revealed four major gene expression patterns, highlighting NEL-dependent regulatory programs (Fig 4F and S5 Fig). Cluster 1 comprised genes with reduced expression specifically in rIpaH7.8-treated cells compared to all other conditions. This cluster was enriched for genes involved in extracellular matrix remodeling, adhesion, and immune signaling, including *CD74*, *SELPLG*, *ANXA1*, and *ADAMTS10* (Fig 4F and S5 Fig). In contrast, Cluster 2 contained genes selectively higher expressed upon rIpaH7.8 treatment. This group included inflammatory mediators and immune regulatory genes such as *IL10*, *EREG*, *PLAUR*, and *ABCA1*, indicating a partial activation or rewiring of inflammatory signaling. Cluster 3 represented genes with reduced expression in the presence of catalytically active rIpaH7.8 but remained elevated in cells treated with the inactive mutant or LPS alone. This pattern indicates a NEL-dependent repression effect (Fig 4F and S5 Fig). Finally, Cluster 4 comprised genes with elevated expression in response to the catalytically inactive rIpaH7.8ΔC, but almost unchanged expression in the presence of active rIpaH7.8 or LPS. This cluster contained classical interferon-stimulated genes and innate immune effectors, including *IFIT1*, *PARP10*, *MAP2K2*, *TRIM22*, and *GSDMD*, highlighting a potential NEL-dependent inhibition of inflammatory and antimicrobial transcriptional programs (Fig 4F and S5 Fig). Interestingly, in contrast to their upregulation in response to LPS, the genes neutrophil cytosolic factor 1 (*NCF1*) in Cluster 4 and its paralog *NCF1b* in Cluster 3 were both downregulated only upon rIpaH7.8 treatment in LPS-stimulated monocytes (Fig 4F and S5 Fig), indicating a reciprocal regulation and thus a downregulation of a pro-inflammatory stimulus. NCF1, also known as p47^phox^, forms a subunit of the NADPH-oxidase-complex NOX2, which catalyzes the production of microbicidal reactive oxygen species (ROS) in phagocytes [30].

**Figure 4.**
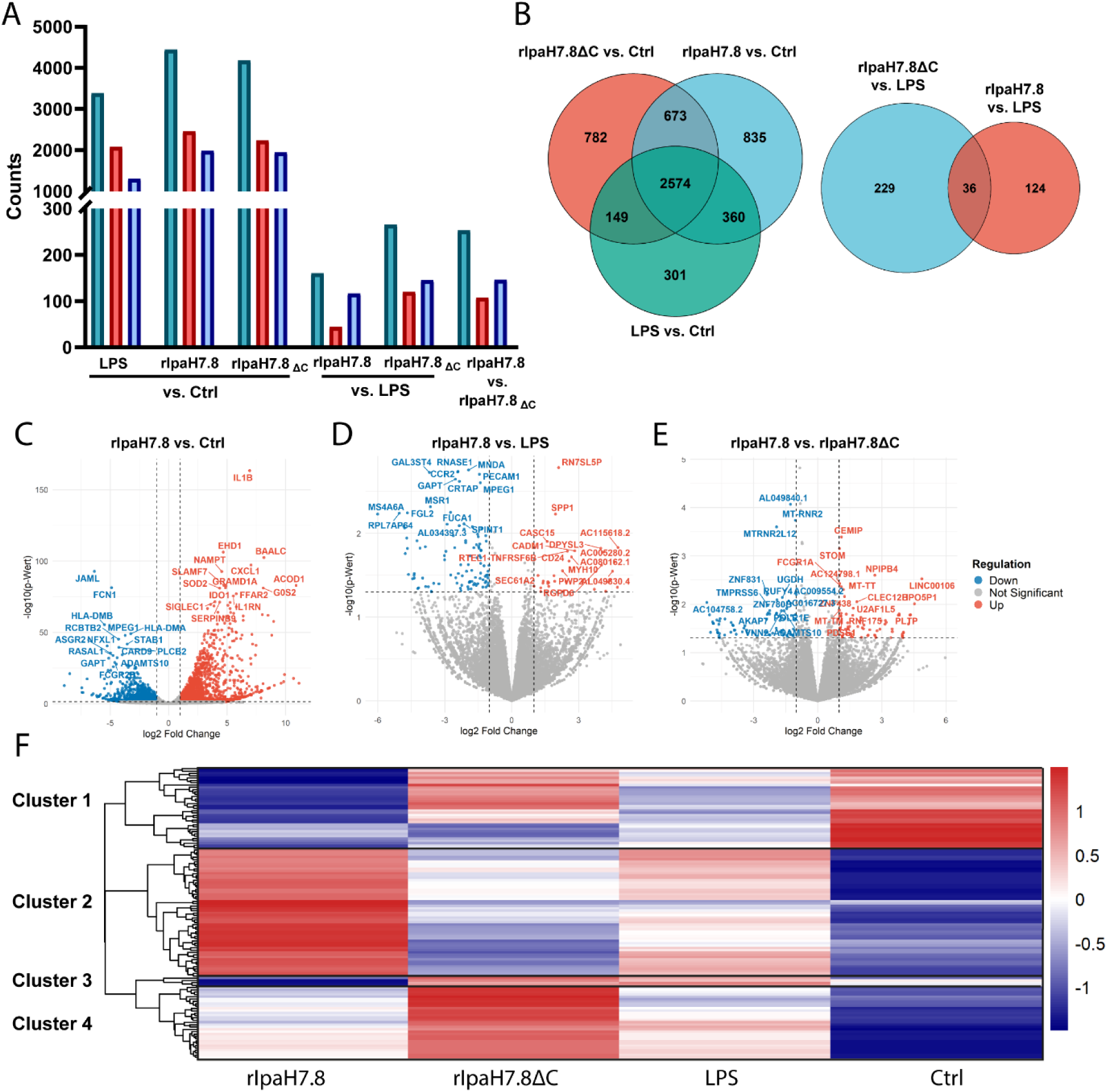
RNA-seq analysis of LPS-stimulated human monocytes treated with rIpaH7.8 or rIpaH7.8ΔC. Primary human monocytes were incubated for 6 h with recombinant rIpaH7.8 or rIpaH7.8ΔC in the presence of LPS; LPS-only and medium-treated cells served as controls. Bulk poly(A)-enriched RNA-seq was performed using three biological replicates per condition, each consisting of pooled RNA from three independent donors. (A) Total number of significantly differentially expressed genes (DEGs; p ≤ 0.05) per comparison. (B) Venn diagrams showing shared and unique DEGs (p ≤ 0.05). Complete DEG lists are provided in Supplementary Data 1. (C-E) Volcano plots depicting differential gene expression in LPS-stimulated monocytes treated with rIpaH7.8 compared to unstimulated controls (C), LPS-only treatment (D), or rIpaH7.8ΔC (E). Significantly up- and downregulated genes (p ≤ 0.05) are shown in red and blue, respectively; the top 30 regulated genes are annotated. (F) Heatmap of rIpaH7.8-regulated genes in LPS-stimulated monocytes relative to rIpaH7.8ΔC, LPS-only, and unstimulated conditions, based on the filtering strategy shown in Supplementary Fig. 4C. Rows represent genes and columns treatment conditions; values are Z-score normalized per gene, with red indicating higher and blue lower relative expression. Unsupervised clustering indicates four major gene expression patterns.

### Functional enrichment links cluster-specific transcriptional programs to immune signaling and vesicular trafficking pathways

To further interpret the functional significance of the cluster-specific transcriptional patterns observed in rIpaH7.8-treated monocytes, we performed Gene Ontology (GO) enrichment analysis of the overlapping NEL-dependent differentially expressed genes identified in the rIpaH7.8 versus rIpaH7.8ΔC comparison under LPS stimulation (S4C Fig). The most significantly enriched biological processes, cellular components, and molecular functions were visualized using cnetplots (Fig 5). Consistent with the transcriptional clustering, several enriched biological processes mapped to genes predominantly contained in Cluster 1 and Cluster 2 (Fig 4F and S5 Fig), reflecting immune regulatory programs. Pathways associated with B-cell activation, differentiation, and proliferation were significantly enriched and included genes such as *INPP5D*, *DDR1*, *TNFRSF21*, *MMP14*, and *CD74* (Fig 5A). These genes, many of which are suppressed in Cluster 1 or induced in Cluster 2, indicate that rIpaH7.8 modulates immune communication networks rather than uniformly repressing them. Notably, *IL10*, primarily associated with Cluster 2, formed a central hub linking immune regulation with wound healing and neuronal signaling. Similarly, *CD74,* encoding a transmembrane glycoprotein, connected immune activation pathways with gene networks related to viral life cycle regulation, suggesting convergence of antibacterial and antiviral responses. In line with the kinase and clustering analyses, several enriched GO terms related to phosphorylation and signaling regulation, such as “positive regulation of transferase activity” and “positive regulation of kinase activity”, were identified. These terms included genes distributed across clusters, including upregulated *CD24*, *EREG*, *CEMIP*, and *LCP2* (Cluster 2), as well as downregulated regulators such as *CHTF18* and *HSP90AA1* (Cluster 1) (Fig 5A). Analysis of the cellular component category further linked specific transcriptional clusters to defined subcellular processes. Terms such as “protein complex involved in cell adhesion” and “focal adhesion” involved genes including *ITGA5*, *ITGB1*, *PLAU*, and *PLAUR*, which were primarily enriched in Cluster 2, consistent with activation of adhesion and remodeling pathways (Fig 5B). In contrast, enrichment of the term “basal part of cell”, including transporter genes such as *SLC11A1*, *SLC1A3*, and *SLC22A4*, as well as *ABCA1*, reflects broader metabolic and membrane-associated adaptations. Interestingly, multiple enriched terms associated with vesicular trafficking and endocytosis mapped predominantly to genes within Cluster 3 (Fig 4F and S5 Fig), which exhibited NEL-dependent repression (Fig 5B). This group included downregulated genes such as *LMAN2L*, *YIPF6*, and *NCF1*, as well as upregulated genes such as *EREG*, *CEMIP*, and *SCARF1*. The selective modulation of these trafficking-associated genes suggests that rIpaH7.8 interferes with intracellular transport and receptor dynamics, consistent with a role in reshaping host cell signaling platforms. Within the molecular function category, enriched terms included PDZ domain binding and immune receptor activity (Fig 5C). These were driven by genes such as *CCR5*, *CCRL1*, and *IFNGR1*, which were predominantly associated with Cluster 2, indicating selective enhancement or rewiring of receptor-mediated signaling pathways rather than global suppression. In addition to these NEL-dependent effects, we identified a subset of genes that were similarly regulated in rIpaH7.8- and rIpaH7.8ΔC-treated monocytes compared to LPS-only controls, indicating NEL-independent transcriptional responses (Fig 4B, S4C Fig). These genes did not map strongly to a single cluster but were enriched for pathways related to MHC class II antigen presentation and immune effector regulation, including *HLA-DPA1*, *HLA-DPB1*, *HLA-DMB*, and *MPEG1* (S6 Fig A). Enriched cellular components included endocytic and lysosomal membranes, and molecular function terms highlighted MHC class II complex activity and pattern recognition receptor signaling, including downregulation of *CLEC7A* (S6 Fig B-D). Taken together, integration of clustering and functional enrichment analyses reveals that rIpaH7.8 induces a pronounced transcriptional reprogramming. Clusters 1 and 4 predominantly reflect suppression of immune effector and inflammatory pathways, Cluster 2 captures induced regulatory and remodeling responses, and Cluster 3 highlights NEL-dependent interference with vesicular trafficking and oxidative burst machinery. Notably, terms linked to phosphorylation and kinase activity included genes distributed across clusters, reinforcing the notion that transcriptional reprogramming by rIpaH7.8 is closely coupled to alterations in host signaling networks.

**Figure 5.**
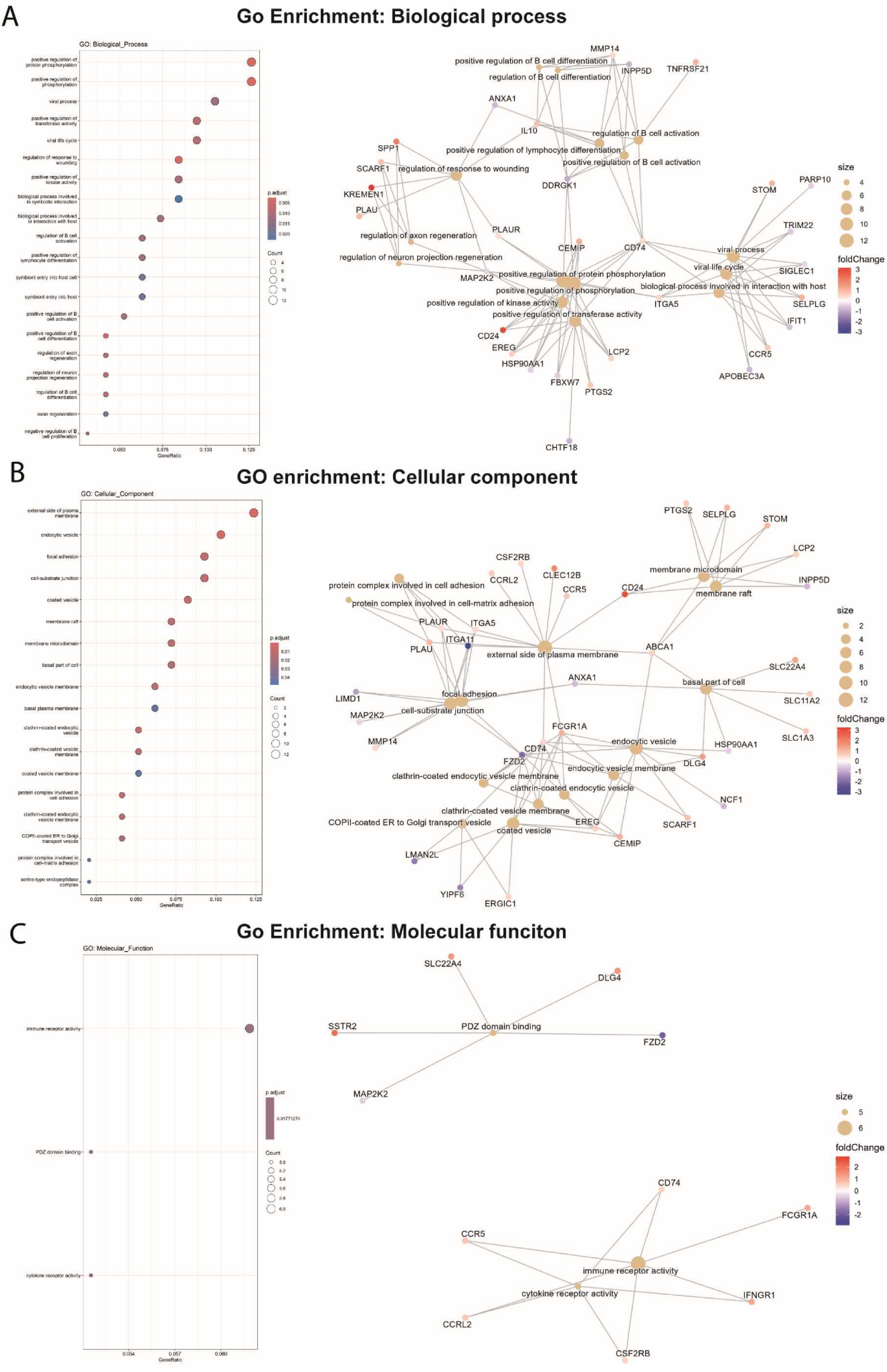
Gene Ontology (GO) enrichment analysis of NEL-dependent overlapping DEGs. GO enrichment of NEL-dependent differentially expressed genes is shown for (A) Biological Processes, (B) Cellular Components, and (C) Molecular Functions. Left panels: Bubble plots showing the top enriched GO terms. The x-axis indicates the GeneRatio, bubble size reflects the number of genes per term (Count), and color intensity represents enrichment significance (adjusted p-value). Right panels: cnet plots visualizing gene GO term associations. Central nodes represent GO terms and peripheral nodes represent genes. GO term node size corresponds to the number of associated genes, while gene node color indicates log₂ fold change (blue, downregulated; white, unchanged; red, upregulated).

### IpaH7.8 catalytically reprograms the host kinome by counteracting LPS-induced kinase responses in primary human monocytes

Given the transcriptional evidence pointing toward IpaH7.8-driven modulation of phosphorylation-related signaling (Fig 4 and 5, S4 Fig and S5), we performed an unbiased kinome activity analysis to functionally validate the predicted interference with kinase activity. Primary human monocytes were stimulated with LPS and incubated with either full-length and catalytically active rIpaH7.8 or its catalytically inactive mutant (rIpaH7.8ΔC). Unstimulated and LPS-only treated cells served as controls. Kinase activity was assessed using the PamGene platform, which monitors phosphorylation of peptide substrates representing known kinase targets, distinguishing between protein tyrosine kinases (PTKs) and serine/threonine kinases (STKs) [31]. The phosphoproteomic analysis on the peptide level revealed a distinct impact of rIpaH7.8 on host signaling pathways (Fig 6A). Although LPS stimulation alone induced changes in several PKT-associated phosphorylation events, such as reduced phosphorylation of TNNT1 (Tyr9), STAT2 (Tyr690), and RBL1 (Tyr805/813), rIpaH7.8 partially counteracted these LPS-driven effects. Conversely, LPS-triggered hyperphosphorylation of pyruvate dehydrogenase E1 (ODPAT, Tyr299) was markedly diminished in the presence of catalytically active rIpaH7.8, but not in ΔC mutant-treated cells (S8A Fig, PTK peptides). Analysis of STK-associated substrates further revealed a cluster of peptides showing a pronounced decrease in phosphorylation specifically upon rIpaH7.8 treatment (S8B Fig, STK peptides). This cluster included STMN2 (Ser97), MYPC3 (Ser269/275, Thr274), and CREB1 (Ser129/133), indicating broader interference with transcriptional regulatory networks. As shown in Figure 6A, direct comparison between rIpaH7.8 and rIpaH7.8ΔC identified a distinct subset of significantly altered phospho-sites. Among the most prominently affected targets were peptides derived from VASP, which displayed consistently reduced phosphorylation in rIpaH7.8-treated cells, suggesting interference with cytoskeletal remodeling and vesicle-associated processes previously implicated in bacterial immune evasion [42,43]. Additional regulated phospho-targets included peptides involved in signaling, migration, and cellular stress responses, including NCF1 (Ser328), which also emerged as downregulated in the transcriptomic analysis (S1 Tab). A complete list of differentially phosphorylated peptides is provided in Supplemental Table 2. Upstream kinase analysis (UKA) based on global peptide phosphorylation profiles underlined a regulatory signature induced by rIpaH7.8. Notably, catalytically active rIpaH7.8 exhibited lower PTK activity and modulated STK activity in a pattern not observed with the ΔC mutant, indicating a catalytic activity-dependent reprogramming of host kinase signaling (Fig 6B, S7 Fig A-E). This differential effect was further supported by direct comparison of rIpaH7.8 and rIpaH7.8ΔC, which uncovered a wide range of differentially active kinases (S9A Fig). To visualize the broader kinase landscape affected by IpaH7.8, modulated kinases were mapped onto the human kinome tree (S10 Fig). Affected kinases clustered within multiple families, including AGC, CMGC, and TKL groups, with node size and color indicating specificity and activity score. Detailed changes in kinase activity were further explored using dot-plot heatmaps. Detailed PTK activity profiles (Fig 6C) demonstrated that rIpaH7.8 effectively counteracted LPS-driven downregulation of multiple kinases, including FES, PTK2B, and TXK, completely or partially restoring their activity relative to cells treated with LPS alone or with LPS and the catalytically inactive variant (ΔC). In contrast, treatment with rIpaH7.8ΔC generally mirrored the LPS-induced PTK suppression, indicating that these effects depend on active NEL catalysis. In several cases, such as PTK7, CSK, EPHA10, and FES, rIpaH7.8 even induced a relative upregulation compared with both LPS-only and LPS + ΔC-treated cells. Of note, EPHA1 was uniquely upregulated by rIpaH7.8 independent of LPS stimulation (Fig 6C). Analysis of STK activity (Fig 6D) showed a similarly broad LPS-dependent activation pattern across CDK, CDLK, CHEK1, and ROCK1 families, consistent across all LPS-containing treatments (S9B Fig). However, treatment with rIpaH7.8 produced a distinct catalytic signature, most prominently reflected in the downregulation of CAMK2 family kinases, relative to both control and LPS or ΔC. In contrast, RPS6KL1 emerged as the only STK significantly upregulated specifically by catalytically active rIpaH7.8. Several central STKs, including PRKACA, PRKCA, SGK1/2, and MYLK3, were attenuated across conditions, consistent with the phosphopeptide-based findings (Fig 6D). Consistent with the RNA-seq analysis (S4C Fig), an intersection of rIpaH7.8-affected kinases (both PTK and STK) was performed to draw a robust conclusion from the collected data. In total, 21 kinases were retained for downstream interpretation (S8C Fig, PTK peptides) and are listed as NEL-dependent (S3 Tab) or NEL-independent (S4 Tab). Thirteen kinases were consistently regulated in all three contrasts (S9C Fig), and fourteen additional kinases overlapped between rIpaH7.8 vs. LPS and rIpaH7.8 vs. ΔC comparisons. Three kinases (CDK13, PRKX, CAMK2D) showed shared regulation in both rIpaH7.8-and ΔC-treated cells. To visualize the signaling architecture influenced by rIpaH7.8, these kinases were integrated with relevant phospho-targets and known IpaH7.8 interaction partners (GSDMD, GSDMB, NLRP1, GLMN, Rab13) in a STRING network (Fig 7A). The resulting network revealed a highly interconnected regulatory core. PKCδ (PRKCD) emerged as a central node, linking kinase modulation directly to NCF1, which was consistently downregulated in the RNA-seq and less phosphorylated in the PamGene datasets. Similarly, PKAα (PRKACA) formed a major hub connecting to its substrate VASP, whose phosphorylation was significantly reduced, and to Rab13, which has previously been shown to accumulate at the *Shigella-*containing vacuole in an IpaH7.8-dependent manner [13]. To contextualize these findings within host-pathway biology, an integrated KEGG pathway analysis was performed that incorporated all rIpaH7.8-affected kinases, all RNA-seq DEGs, and additional IpaH7.8-associated proteins (GSDMD, GSDME, GLMN, Rab13). In this regard, the top 10 enriched pathways are shown in Figure 7B. Among them, ErbB signaling emerged as the most prominent pathway, reflecting broad modulation of receptor tyrosine kinase signaling. Several immune- and infection-related pathways were also enriched, including Fcγ receptor-mediated phagocytosis, C-type lectin receptor signaling, leukocyte transendothelial migration, NOD-like receptor signaling, and pathways associated with *Yersinia* infection and Shigellosis (Fig 7C). Additionally, the analysis highlighted multiple canonical immune-regulatory cascades, PI3K-Akt, MAPK, mTOR, and HIF-1 signaling, as well as pathways linked to inflammatory modulators (TRP channel regulation) and host cell survival/proliferation (Fig 7B; neurotrophin signaling, glioma pathways, circadian entrainment). Taken together, the network and pathway analyses demonstrate that rIpaH7.8 shapes a highly interconnected kinase-gene signaling landscape that integrates immune regulation, cytoskeletal remodeling, and vesicular trafficking in primary monocytes, thereby revealing coordinated host pathways potentially exploited during *Shigella* infection.

**Figure 6.**
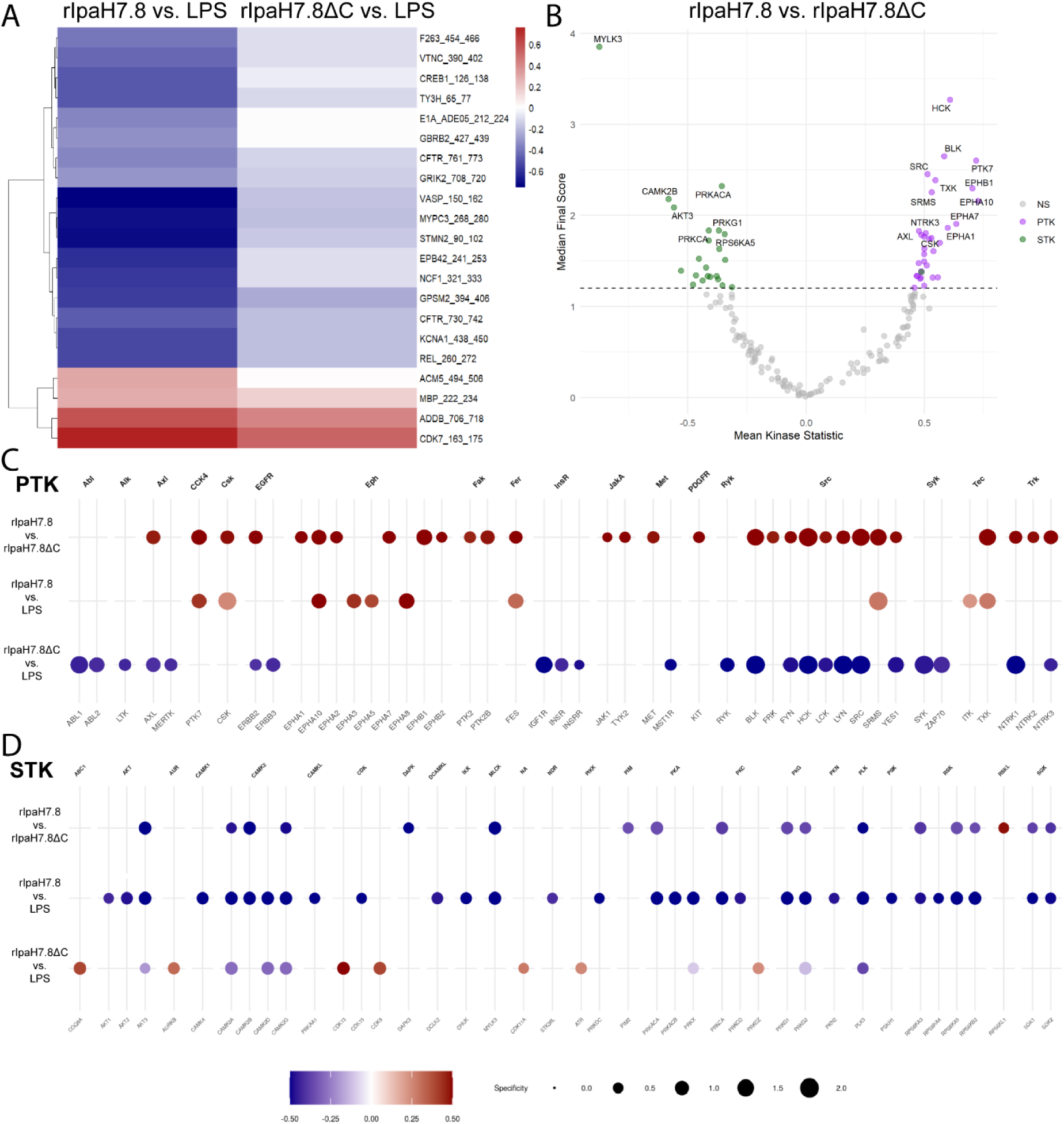
Kinome profiling of LPS-stimulated human monocytes treated with rIpaH7.8 or IpaH7.8ΔC. (A) Heatmap of significantly regulated phosphorylation sites identified by PamChip analysis in LPS-stimulated primary human monocytes treated with rIpaH7.8 compared to the catalytically inactive mutant rIpaH7.8ΔC. Rows represent individual peptide substrates. (B) Volcano plots of inferred upstream kinase activity based on peptide phosphorylation changes. Comparisons shown are LPS vs. control, LPS + rIpaH7.8 vs. control, and LPS + rIpaH7.8ΔC vs. control. Each dot represents one peptide; significantly regulated peptides (median final score > 1.2) are highlighted (PTK, purple; STK, green). (C, D) Kinase activity maps for protein tyrosine kinases (PTK; C) and serine/threonine kinases (STK; D). Color indicates relative changes in inferred kinase activity (blue, decreased; white, unchanged; red, increased), and node size reflects substrate specificity.

**Figure 7.**
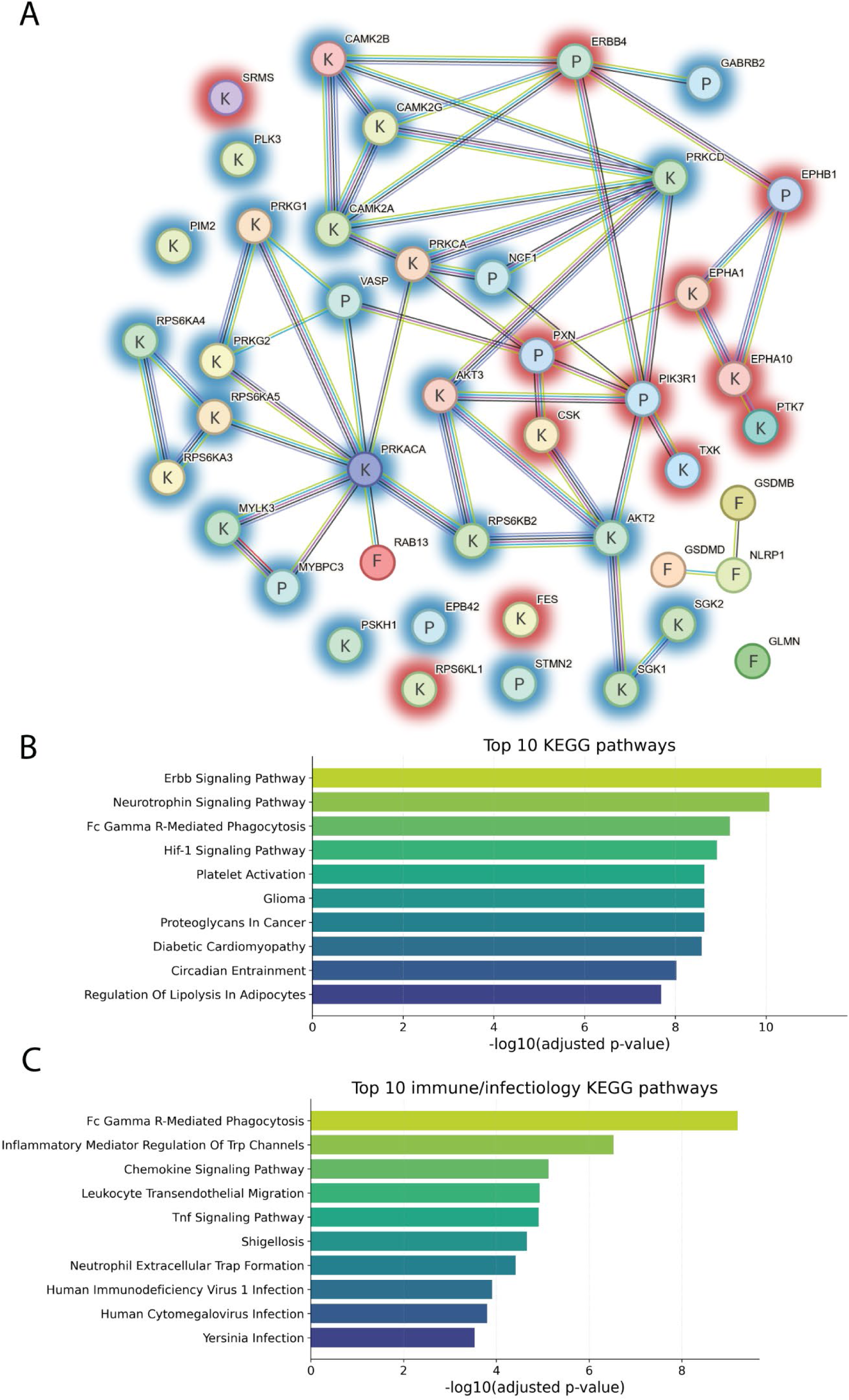
rIpaH7.8-centered signaling networkand enriched host pathways. (A) STRING network analysis integrating differentially phosphorylated peptides (Table S1), regulated kinases (Tables S3 and S4), and IpaH7.8-associated proteins (GLMN, GSDMD, GSDMB, NLRP1, Rab13; underlined). Edge colors indicate STRING evidence types: curated databases (red), experimentally determined interactions (blue), gene neighborhood (green), co-expression (black), protein homology (purple), text mining (light blue), and interactions without assigned evidence (grey). Factors (F), Kinase (K), and Peptid (P) are shown, Halos indicated up- (red) or down- (blue) regulation. (B-C) KEGG pathway enrichment analysis based on genes and kinases significantly modulated in transcriptomic and PamGene kinome profiling datasets. Shown are the top 10 enriched pathways (B) ranked by statistical significance (-log10 adjusted P value, Benjamini-Hochberg FDR) and the top 10 infection- and immunology-related KEGG pathways (C). Bar length indicates enrichment significance.

### rIpaH7.8 attenuates NOX2-associated kinase signaling and reactive oxygen species production in LPS-stimulated human monocytes

Integration of kinase activity profiling with STRING network and KEGG pathway analyses revealed that rIpaH7.8 preferentially targets central signaling hubs linked to inflammatory and antimicrobial effector pathways. Within the rIpaH7.8-centered STRING network (Fig 7A), several kinases inferred to be attenuated in the upstream kinase analysis clustered around components of the NOX2 complex, including NCF1/p47^phox^ (Fig 8A). Among these, protein kinase C isoforms (PRKCα and PRKCδ) and protein kinase A catalytic subunit α (PRKACA) emerged as key connecting nodes linking kinase signaling to cytoskeletal regulation, vesicular trafficking, and immune effector functions. Although PRKCα displayed very low transcript levels in human monocytes (FPKM < 1), PRKCδ was abundantly expressed (FPKM > 20) and transcriptionally stable across all conditions, indicating post-transcriptional or activity-based regulation (RNA seq data GSE324547). Consistently, phosphorylation of NCF1-derived peptides, including regulatory serine residues targeted by PKC family members, were reduced in rIpaH7.8-treated cells (Fig 6A and 8A). Additional candidate kinases included AKT3, PKCγ, and PRKACA, the latter of which occupies a central position within the STRING network. RNA-seq analysis revealed significant downregulation of PRKACA upon LPS stimulation, which was further enhanced by rIpaH7.8 treatment, consistent with reduced inferred kinase activity. KEGG enrichment analysis of integrated transcriptomic and kinome datasets highlighted Fcγ receptor-mediated phagocytosis, chemokine signaling, TRP channel regulation, and PI3K-Akt signaling (Fig 7B), all of which are functionally linked to PKC/PKA activity and NOX2-dependent ROS production. In parallel, multiple NOX2 complex components (NCF1/p47^phox^, p40^phox^, and gp91^phox^) were transcriptionally downregulated, indicating a coordinated suppression of the oxidative burst machinery (Fig 8A). To functionally validate the network- and pathway-based prediction of impaired NOX2 activity, intracellular reactive oxygen species (ROS) production was quantified using a DCFDA-based cellular ROS assay (Fig 8B). Consistent with the kinase and transcriptomic data, LPS and PMA stimulation increased ROS production in primary human monocytes, whereas treatment with rIpaH7.8 significantly reduced LPS-induced ROS levels (Fig 8B). In contrast, the catalytically inactive rIpaH7.8ΔC variant had no significant effect on ROS production. Together, these data demonstrate that rIpaH7.8-mediated reprogramming of host kinase signaling translates into a measurable suppression of NOX2-dependent oxidative burst responses in human monocytes.

**Figure 8:**
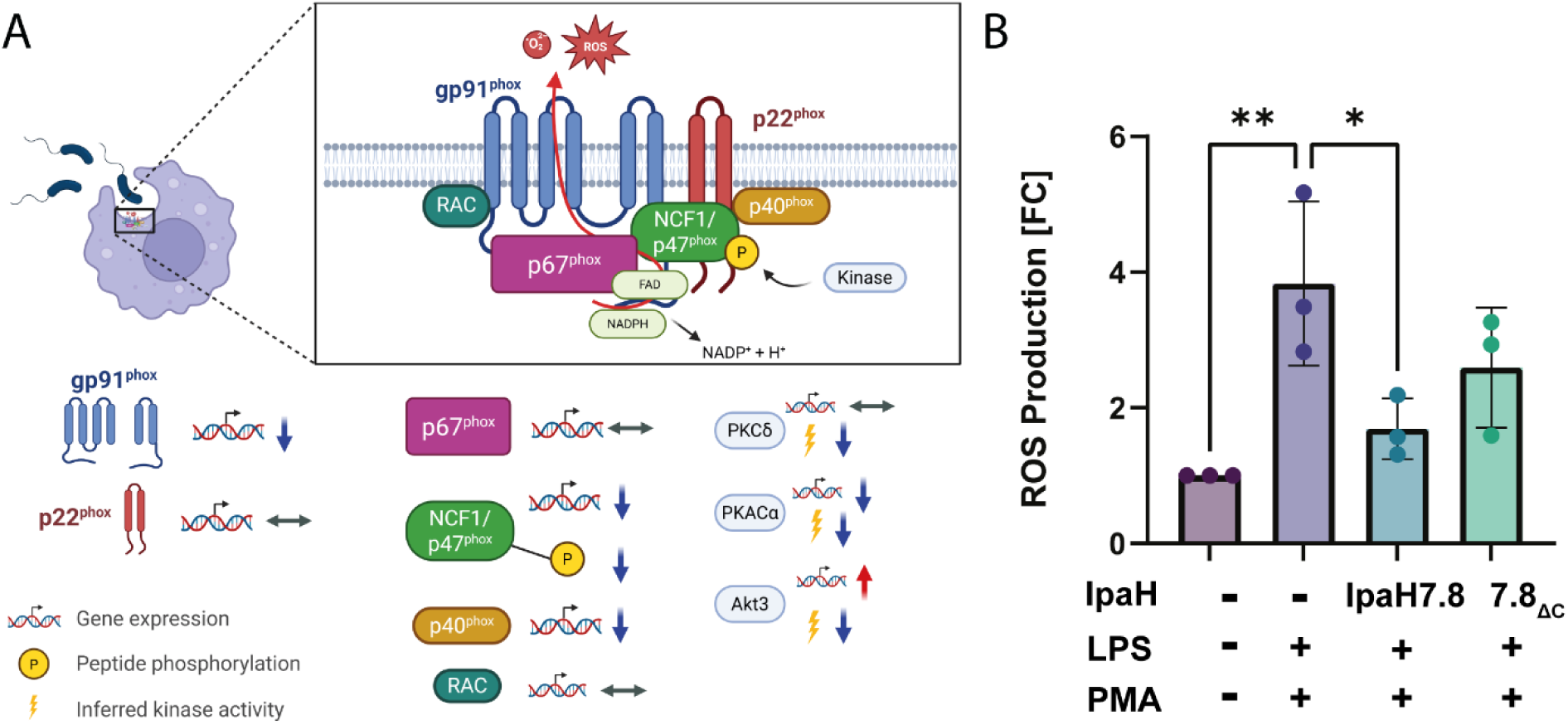
rIpaH7.8 reduces ROS production in LPS-stimulated human monocytes. (A) Structure of the activated NOX2 complex, comprising the integral membrane proteins gp91^phox^, p22^phox^ and associated components p40^phox^, NCF1/p47^phox^, p67^phox^ and Rac1 or Rac2 assembled at the plasma- or phagocytic membrane (upper panel). Lower panel indicates regulation of NOX2 complex proteins upon treatment of primary human monocytes with LPS+rIpaH7.8 (1 mg/ml; 1.6 μM) on transcriptional, phosphorylation or kinase-activity level. Scheme adapted from Bode et al. 2023 [67]. (B) Cellular reactive oxygen species (ROS) production in LPS-stimulated primary human monocytes following 2h PMA stimulation in the presence or absence of rIpaH7.8 or rIpaH7.8ΔC. One-way ANOVA with Tukey’s post hoc test (ns, non significant, **p < 0.01, *p < 0.05).

## Discussion

In this study, we demonstrate that the *Shigella* effector IpaH7.8 possesses an intrinsic, T3SS-independent capacity to enter eukaryotic cells, expanding the functional repertoire of cell-penetrating LPX effectors. Recombinant IpaH7.8 (rIpaH7.8) entered host cells via macropinocytosis and lipid raft-dependent endocytosis, in a temperature-dependent manner consistent with an energy-requiring process (Fig 2, S1 Fig and S2). Chemical inhibition of macropinocytosis and actin dynamics significantly reduced uptake, highlighting the contribution of cytoskeletal remodeling. Similar to other LPX effectors (IpaH9.8, SspH1, YopM) [14,17,24,32], the N-terminal α-helical region of IpaH7.8 functions as a protein transduction domain (PTD) enabling autonomous internalization. Following uptake, rIpaH7.8 localized to endo-lysosomal compartments but also reached the cytosol, as indicated by pH-sensitive fluorophore quenching and by polyubiquitination of cytosolic GSDMD, which required the intact NEL domain (Fig 2 and Fig 3). This confirms that a fraction of internalized protein escapes endosomal degradation. While the precise escape mechanism remains unknown, the PTD exhibits pH-dependent modulation of net charge, becoming slightly more positive in acidic compartments (Fig 1C), a feature shared with other pH-sensitive anionic CPPs such as Sap(E) [33]. As described for Sap(E) and other negatively charged CPPs, uptake appears to rely on a multifactorial mechanism combining endocytosis, macropinocytosis, and local membrane interactions [34,35]. These similarities suggest that LPX PTDs form a distinct class of anionic, pH-responsive CPEs optimized for cytosolic delivery during infection. Whether T3SS-independent uptake contributes to *Shigella* virulence *in vivo* remains unresolved. Nonetheless, CPE-mediated delivery could theoretically enable effectors to act on bystander or distal host cells not directly contacted by bacteria, an advantage unavailable to classical T3SS injection. Using this T3SS-independent delivery approach, we show that the catalytic NEL domain remains fully functional in host cells, mediating polyubiquitination of endogenous GSDMD. In contrast, intracellular ubiquitination of GLMN, which was previously proposed as a key IpaH7.8 substrate in mice [3], was not detectable, consistent with species-specific differences reported by Luchetti *et al.* 2021 [9]. Functionally, GSDMD modification by rIpaH7.8 resulted in a marked reduction of IL-1β secretion following LPS priming and Nigericin activation in both THP-1 macrophages and primary monocytes (Fig 3, S3 Fig). rIpaH7.8 did not impact upstream inflammasome assembly, as ASC speck formation remained unaffected (Fig 3F), supporting a model in which IpaH7.8 acts downstream of inflammasome activation, consistent with Luchetti *et al.* [9].

To dissect broader immunomodulatory effects, we performed RNA-seq on LPS-stimulated monocytes treated with either full-length rIpaH7.8 or the catalytic deletion variant rIpaH7.8ΔC (Fig 4). This experimental design allowed us to distinguish between transcriptional effects mediated by the cell-penetrating properties and LRR target-binding domain (shared by both constructs, Fig 1A, Fig 2B), and those specifically driven by the catalytic activity of the NEL ubiquitin ligase domain, which is deleted in the ΔC variant. The comparison of full-length IpaH7.8 to both untreated cells and its catalytically inactive ΔC variant revealed a distinct transcriptional signature attributable to its enzymatic activity (NEL-dependent, Fig 4 and S4 Fig). Notably, the transcriptional landscape induced by rIpaH7.8 was not uniformly altered but segregated into distinct gene clusters, reflecting coordinated and functionally separable host responses. Genes grouped in Cluster 1 and Cluster 4 were predominantly suppressed and included key mediators of innate immune effector functions, most prominently components of the NOX2 complex such as NCF1, consistent with the observed attenuation of ROS production. In contrast, Cluster 2 comprised genes that were selectively upregulated, including regulatory and immunomodulatory factors such as IL10 and CD24, suggesting that rIpaH7.8 does not simply silence host responses but actively reshapes them toward a more controlled or anti-inflammatory state. A third group of genes (Cluster 3) displayed a characteristic NEL-dependent regulation pattern, being suppressed specifically by catalytically active rIpaH7.8 but not by the ΔC variant. This cluster was enriched for genes involved in vesicular trafficking and endosomal dynamics linking the enzymatic activity of IpaH7.8 to interference with intracellular transport processes and antimicrobial effector mobilization. Thus, rIpaH7.8 induces a defined, cluster-resolved transcriptional program that functionally separates immune suppression, regulatory rewiring, and vesicle-associated interference into distinct but interconnected modules. In this regard, Gene Ontology analysis of NEL-dependent DEGs revealed strong enrichment for pathways linked to kinase regulation, phosphorylation signaling, and immune system processes, indicating that rIpaH7.8 reshapes host signaling both through direct protein modification and by altering transcriptional programs. Among the most prominent categories were endocytic vesicles and membrane microdomains, supporting the notion that rIpaH7.8 is internalized autonomously through energy-dependent endocytosis (Fig 5). Within this cluster, rIpaH7.8 strongly downregulated key regulators of vesicle trafficking, including LMAN2L and YIPF6, both of which are essential for ER-Golgi transport and innate immune functions [36,37]. Additionally, IpaH7.8-regulated candidates, such as RUFY4 and TBC1D2 (Armus), implicate the effector in modulating late endosomal maturation and autophagy [38,39]. Notably, Armus also acts downstream of Rac1, which is activated by the *Shigella* effector IpgB1, raising the possibility that rIpaH7.8 supports IpgB1-dependent epithelial invasion and vacuolar escape [40,41]. A second major cluster involves components of the NOX2 complex, including NCF1, all downregulated by rIpaH7.8 (Fig 8A). Such coordinated suppression of ROS-generating machinery, together with the known degradation of GSDMB by IpaH7.8 [10], suggests that the effector might employs a multilayered strategy to inhibit both oxidative and non-oxidative killing pathways within phagocytes. Within the enriched GO category “positive regulation of kinase activity,” CD24 emerged as one of the most robustly upregulated genes (Fig 5A). The CD24-Siglec-10 axis dampens DAMP-induced inflammation [42] and can function as an anti-phagocytic (“don’t eat me”) signal [43]. rIpaH7.8-mediated CD24 upregulation may therefore provide *Shigella* with an advantage by suppressing macrophage responsiveness while enhancing epithelial motility pathways relevant to infection. Intersecting DEGs shared between rIpaH7.8 and rIpaH7.8ΔC identified 36 NEL-independent genes (Fig 4B), reflecting effects driven by protein uptake or LRR-mediated target binding (Fig 3). The strongest signature was a coordinated downregulation of MHC class II genes, including HLA-DPA1, HLA-DPB1, HLA-DMB, and HLA-DRB1. This pattern suggests modulation of the master regulator CIITA, reminiscent of pathogen-driven suppression of antigen presentation [44,45]. Importantly, despite efficient protein delivery, neither rIpaH7.8 nor rIpaH7.8ΔC induced signatures of endosomal rupture or lysosomal stress, unlike some cationic CPPs [46]. This supports functional translocation without cytotoxicity and indicates that the LRR domain alone can modulate immune gene networks without provoking global stress responses. To functionally validate and link the transcriptomic evidence with phosphorylation-related pathway modulation, we performed kinome profiling on LPS-stimulated primary human monocytes treated with either catalytically active rIpaH7.8 or its inactive variant. This analysis revealed that IpaH7.8 broadly reprograms kinase signaling, rather than acting on a single linear pathway (Fig 6). Strikingly, several protein tyrosine kinases (PTKs) suppressed by LPS, consistent with monocyte adaptation to sustained TLR4 stimulation, were partially restored in the presence of catalytically active rIpaH7.8 (S 8 Fig & S9). These effects were absent in cells treated with the inactive mutant, indicating a NEL-dependent interference with LPS-driven kinase attenuation. Although some predicted kinases (e.g., EPH and TRK families) were not detected in RNA-seq datasets, several candidates, including FES, CSK, PTK2B, and TXK, were well expressed, supporting a genuine regulatory effect. Among the most consistently affected nodes were CaMKII isoforms, which displayed reduced inferred activity in rIpaH7.8-treated cells (Fig 6; S9 Fig). CaMKIIγ/δ function as key amplifiers of TLR-induced NF-κB, IRF3, and MAPK signaling, thereby boosting inflammasome priming and cytokine production. Their attenuation by IpaH7.8 may therefore represent an additional layer of immune dampening, complementary to the effector’s direct modification of GSDMD. Notably, CaMKII transcripts showed only minimal changes, suggesting that IpaH7.8 affects CaMKII function through post-transcriptional or network-level mechanisms. Similarly, Akt2 and Akt3 emerged as central serine/threonine kinases affected by rIpaH7.8. Although Akt transcript levels remained stable, the upstream kinase analysis indicated a reduction in inferred activity specifically in the presence of catalytically active IpaH7.8. Beyond their roles in metabolic and polarization states of macrophages, Akt3 also phosphorylates NCF1/p47^phox^, a key component of the NOX2 complex involved in phagosomal ROS production [47]. Consistent with this, the PamGene data revealed a significant reduction of NCF1 Ser328 phosphorylation only in rIpaH7.8-treated monocytes (Fig 6C; S2 Tab), implicating altered Akt and PKC signaling as a mechanistic basis for impaired oxidative burst induction. Indeed, PKCδ, a master regulator of inflammatory signaling and a known NCF1/p47phox kinase, appeared as another major node attenuated by rIpaH7.8. PKCδ activity is central for phosphorylation-dependent assembly of the NOX2 complex and for efficient microbial killing by macrophages [48]. The combined decrease in PKCδ-associated activity, reduced phosphorylation of NCF1, and transcriptional downregulation of *NCF1*, *NCF4*, and *CYBB* in RNA-seq data (Fig 8A; S1 Tab) strongly suggests that rIpaH7.8 suppresses NOX2-mediated ROS production through multi-level interference which translates into a measurable suppression of NOX2-dependent oxidative burst responses in human monocytes (Fig 8B). A similar virulence strategy has been described for *Francisella tularensis*, which inhibits PKC-dependent NCF1 activation to block the oxidative burst [49]. Our findings, therefore, raise the possibility that *Shigella* exploits IpaH7.8 to dampen phagosomal ROS generation during infection. In addition to the NOX2 axis, the kinase network contained a smaller module involving PRKACA and VASP, both of which were previously linked to epithelial barrier integrity and, intriguingly, to the regulation of *Shigella* vacuolar escape [13,50]. Although VASP phosphorylation changes were modest in monocytes, the convergence of PRKACA, Rab13, and VASP within a single signaling module (Fig 7A and S11 Fig A & B) may point to a broader role for IpaH7.8 in modulating cytoskeletal and vesicular trafficking processes beyond inflammasome inhibition. However, this needs to be investigated in future experiments in the context of *Shigella* infection, including relevant mutant strains. Overall, integration of the kinome and transcriptomic datasets reveals that IpaH7.8 not only suppresses inflammasome effector function via direct ubiquitination of GSDMD but also reconfigures phosphorylation networks that govern TLR responses, CaMKII- and Akt-driven inflammatory amplification, and PKC/NOX2-dependent antimicrobial pathways. This multi-layered interference underscores IpaH7.8 as a highly versatile immune modulator that operates simultaneously at transcriptional, post-transcriptional, and post-translational levels. Interestingly, Wang et al. 2023 discovered a third inhibitory strategy, in which IpaH7.8 conducts mono-ubiquitinations of the lysines K177, K190, and K192 of gasdermin B [51]. Future phosphoproteomic mapping and substrate-trapping NEL mutants will be essential to resolve whether the observed kinase alterations arise from direct ubiquitin ligase activity or represent secondary rewiring of host signaling networks.

In conclusion, this study reveals that IpaH7.8 is a dual-function effector that combines autonomous cell entry with catalytic interference in host ubiquitin and phosphorylation networks. It suppresses pyroptotic signaling through direct ubiquitination of GSDMD, counteracts LPS-induced kinase responses, reprograms cytoskeletal signaling by inhibiting PKA-dependent pathways, and reduces NOX2-mediated ROS production. These combined activities may facilitate *Shigella flexneri* vacuolar escape and intercellular spread while attenuating host inflammatory responses. The anti-inflammatory properties of rIpaH7.8, particularly its ability to suppress IL-1β release via GSDMD ubiquitination, highlight its potential for therapeutic exploitation. Recombinant IpaH7.8 or engineered PTD-fusion proteins could serve as targeted immunosuppressive tools in diseases characterized by excessive pyroptosis or IL-1-driven inflammation, such as autoinflammatory syndromes (e.g., familial Mediterranean fever), sepsis, or inflammatory bowel disease (IBD). However, the enzyme’s broad substrate spectrum and its ability to rewire host signaling networks also pose potential risks. Off-target ubiquitination or suppression of protective immune responses could increase susceptibility to infection or oncogenic transformation. Therefore, any therapeutic application would require precise control of delivery, activity, and tissue specificity. Beyond the direct translational potential, a better understanding of how microbial effectors such as IpaH7.8 manipulate host signaling may reveal an extensive natural repertoire of cell-penetrating immunomodulators. This shift in perspective - from viewing effectors solely as virulence factors to recognizing them as molecular tools - was previously articulated as the ‘drugs from bugs’ concept [59–62].

## Material and Methods

### Antibodies

Anti-FLAG M2 (#F1804, Sigma Aldrich), anti-GAPDH (#sc-25778, Santa Cruz), anti-Rab5 (#3547, Cell Signaling Technology), anti-Rab7 (#9367, Cell Signaling Technology), anti-CD63 (#sc-15363, Santa Cruz), anti-HA (#2367 Cell Signaling Technology), anti-Ubiquitin (#sc-8017; Santa Cruz,), anti-Gasdermin-D (#97558, Cell Signaling Technology), anti-Glomulin (#PA5-27838; Invitrogen), anti-GFP (#MAB3580; Merck Millipore), anti-transferrin-receptor (#13-6800, invitrogen) HRP-labeled goat anti-mouse IgG (#115-035-003, Dianova), HRP-labeled goat anti-rabbit IgG (#111-035-003, Dianova), Cy2 labeled goat anti-mouse IgG (#115-225-166, Dianova), Cy2 labeled goat anti-rabbit IgG (#111-22-003, Dianova), and rabbit IgG anti-mouse IgG+IgM (#115-225-166, Dianova) antibodies were obtained commercially.

### Plasmids

Plasmids for the subsequent overexpression of the recombinant proteins IpaH7.8-3xFLAG, 2αH-IpaH7.8-GFP-3xFLAG, and GFP-3xFLAG were cloned based on the pET-24b(+) 3xFLAG or the pET-24b(+) vector, respectively [14]. All of the above-named plasmids were generated using a restriction-free cloning approach [56]. The plasmids for the overexpression of recombinant GFP-3xFLAG, YopM-3xFLAG, and 2αH-YopM-GFP-3xFLAG have been previously constructed [14,20]. Plasmids for the overexpression of IpaH7.8ΔN (lacking amino acids 1-57) and IpaH7.8ΔC (lacking amino acids 359 - 519) were constructed by deleting the DNA sequence coding for the indicated amino acids by inverse PCR.

### Expression, purification, and fluorescent labeling of recombinant proteins

*Escherichia coli* Clear Coli® BL21 (Lucigen, Middleton, WI, USA) harboring the respective plasmids was cultured in Lysogeny Broth (LB) medium supplemented with kanamycin (50 µg/mL) at 37°C, with an optical density at 600 nm (OD600) of 0.7. Protein expression was induced by adding 1 mM IPTG and incubating overnight at 16°C. Following bacterial harvesting via centrifugation, cell disruption was performed by sonication on ice for 30 seconds with 30-second breaks, repeated six times (Branson Sonifier 250, level 4, 50% cycle; Branson, Danbury, CT, USA). Isolation and purification of 6xHis-tag fusion proteins was carried out by affinity chromatography using nickel-nitrilotriacetic acid (Ni-NTA) agarose beads (Macherey-Nagel, Düren, Germany), as previously described [15]. The purified proteins were then dialyzed against phosphate-buffered saline (PBS) and concentrated using Centricon centrifugal filters (Merck Millipore, Billerica, MA, USA). For conjugating recombinant proteins with fluorescent dyes, the Cy3 antibody labeling kit (GE Healthcare, Chicago, IL, USA), the Alexa Fluor® 488 antibody labeling kit (Thermo Fisher Scientific, Waltham, MA, USA), the FluoReporter™ FITC protein labeling kit (Thermo Fisher Scientific), or 5-(and 6-)carboxynaphthofluorescein succinimidyl ester mixed isomers (naphthofluorescein, NF) (Thermo Fisher Scientific) were used according to the manufacturer’s protocols.

### Cell culture

HeLa cells were cultured in DMEM with low glucose (1 g/L) (Sigma-Aldrich), supplemented with 10% fetal calf serum (FCS) and 5% non-essential amino acids. HEK293T cells were maintained in DMEM with high glucose (4.5 g/L) (Sigma-Aldrich), supplemented with 10% FCS and 5% non-essential amino acids. THP-1 cells were cultured in RPMI-1640 (Sigma-Aldrich), supplemented with 10% Heat-inactivated FCS (Hi-FCS). All cell lines were kept at 37°C in a 5% CO2 atmosphere, and the growth medium was replaced every 2-3 days. The differentiation of THP-1 cells into THP-1- derived macrophages was achieved by adding 25 ng/mL of phorbol 12-myristate 13-acetate (PMA, BioGems) and subsequently resting the cells. On day 3 after the addition of PMA, cells were gently washed with PBS and rested again until Day 5, when experiments were performed.

### Isolation of primary human monocytes

Primary human monocytes were isolated from leukocyte reduction system (LRS) chambers obtained from fresh platelet apheresis donations of anonymized healthy volunteers after informed consent of the donors according to the regulations of the blood bank of the University Hospital Münster [57]. Briefly, the chambers were drained and subsequently flushed with sterile 1x HBSS to recover all residual cells. The blood-HBSS-mix was carefully layered onto a Pancoll gradient (human Pancoll; density: 1,077g/ml PAN Biotech). After centrifugation at 600 x g for 35 min at room temperature (Swingbucket rotor; break off), peripheral blood mononuclear cells (PBMCs) were collected from the interphase and subjected to several washing steps with HBSS. For further Monocyte enrichment and lymphocyte depletion, the PBMCs were applied to a second density gradient using Percoll-® (Cytiva). After 60 min centrifugation at 540 x g (brake off), the upper monocyte-enriched layer was collected and washed with HBSS, and finally resuspended in RPMI 1640 medium supplemented with 10% Hi-FCS and a cell density of 2 x 10^6^ cells/mL. Monocyte purity (at least 85%) was assessed by flow cytometry. After isolation, the monocytes were rested overnight in Lumox-foil bags and used the next day for downstream applications.

### Fractionation of eukaryotic cells

For separation of cytoplasmic and membrane proteins of HeLa cells, cell fractionation was performed following incubation of the cells with the recombinant proteins (0.8 µM) for 3 h as previously described and analyzed by immunoblotting [20,65–67]. Purity of cytoplasmic and membrane fractions was analyzed using antibodies against soluble cytosolic glyceraldehyde-3-phosphate dehydrogenase (GAPDH) and the membrane marker transferrin receptor (TF-R), respectively.

### Confocal laser scanning microscopy (CLSM)

Cells were cultured in 24-well plates until they reached 70% confluency. Adherent cell lines (HeLa, HEK293T) were directly seeded onto glass coverslips, while suspension cell lines (THP-1) were first allowed to attach to poly-L-lysine-coated coverslips before fixation. Fluorescently labeled proteins were then added to the cells. Following incubation, cells were washed three times with PBS, and residual surface-bound proteins were removed by treatment with an acid buffer (0.2% w/v glycine in PBS, pH 2.0) for 5 min, followed by three additional PBS washes. Cells were fixed using 4% w/v paraformaldehyde (PFA) in PBS and permeabilized with 0.2% w/v Triton X-100. To visualize filamentous actin, cells were stained with tetramethylrhodamine (TRITC)- or fluorescein isothiocyanate isomer I (FITC)-conjugated phalloidin at a 1:500 dilution in PBS for 30 min. DNA was stained with DRAQ5 at a 1:500 dilution in PBS for 30 min or Dapi at a 1:1000 dilution in PBS for 8 min. For additional cellular structure labeling, primary antibodies specific to Rab5 (early endosomes), Rab7 (late endosomes), and CD63 (lysosomes) were used at dilutions of 1:100, 1:50, and 1:600 in PBS, respectively, after blocking with 5% v/v goat serum in PBS for 1 h. Appropriate Cy2-labeled secondary antibodies were applied at a 1:100 dilution in PBS for detection. Cells were washed three times with PBS after each step. The preparations were mounted in DAKO fluorescent mounting medium and analyzed using a Zeiss LSM 510 Meta or a Leica Stellaris 5 confocal laser-scanning microscope, utilizing the appropriate filters.

### Uptake analysis

Uptake kinetics of internalized proteins were monitored using flow cytometry (FACScan flow cytometer; BD Biosciences, San Jose, CA, USA and Cytoflex S, Beckman Coulter). Cellular translocation was analyzed by continuous time-lapse quenched uptake assays, as previously described [16,61]. In brief, Alexa Fluor® 488-/FITC-labeled proteins were added to trypsinized cells for continuous incubation. To quench extracellular fluorescence during measurement, trypan blue (final concentration 0.2%) was used. To evaluate the effect of various endocytic inhibitors, HeLa cells were grown in 6-well plates to 80% confluency. After pre-incubation with inhibitors (200 µM cytochalasin D (Sigma-Aldrich), 19 mM amiloride (Sigma-Aldrich), 30 mM dynasore (Enzo Life Sciences, Farmingdale, NY, USA), 50 mM methyl-β-cyclodextrin (MβCD, Sigma-Aldrich), 16.5 mM nocodazole (Sigma-Aldrich) for 1 h, 25 µg/mL Alexa Fluor® 488-labeled rIpaH7.8 or its deletion variants were added to the cells for an additional 3 hours. Prior to measurement, cells were trypsinized and quenched with trypan blue as described above. For uptake assays involving naphthofluorescein-labeled proteins, HeLa cells were cultured in 6-well plates to 80% confluence. After incubation with the proteins for the specified time periods, cells were washed three times with PBS and treated with acid buffer (0.2% w/v glycine in PBS, pH 2.0) for 5 minutes to remove any remaining membrane-bound proteins. Unless otherwise stated, cells were incubated at 37°C in a 5% CO2 atmosphere. For each sample, fluorescence intensity (GeoMean) was measured in triplicate for 10,000 cells.

### *In vitro* ubiquitination assays

The enzymatic activity of recombinant proteins as E3 ubiquitin ligases was evaluated through *in vitro* ubiquitination assays. These were conducted in 40 µL reaction mixtures containing reaction buffer (25 mM Tris-HCl, pH 7.5; 50 mM NaCl; 5 mM ATP; 10 mM MgCl2; 0.1 mM DTT), 0.5 µg E1, 2 µg E2 (UbcH5b), 2.0 µg HA-Ubiquitin (all purchased from Boston Biochem, Cambridge, MA, USA), with or without 4 µg of the respective protein of interest. The reaction mixtures were incubated at 37°C for 1 hour, and the reactions were terminated by the addition of 4x Laemmli buffer.

### 3xFLAG Pulldown assay

THP-1-derived macrophages were cultured in ten 10 cm dishes and harvested after differentiation by scraping in 300 µl of lysis buffer (50 mM Tris HCl, pH 7.4, with 150 mM NaCl, 1 mM EDTA, and 1% triton X-100) supplemented with 1x Halt™ Protease Inhibitor Cocktail (100x; Thermo Scientific). Anti-FLAG-M2 Magnetic beads (Sigma-Aldrich) were loaded with 120 µg recombinantly expressed and purified rIpaH7.8, rIpaH7.8ΔC, both carrying a C-terminal 3xFLAG-tag (DYKDDDDK). Control beads were either loaded with rGFP-3xFLAG or left unloaded. Bead loading was performed by incubating the beads with the bait proteins for 1 h at room temperature under constant shaking. After loading, the beads were washed twice with TBS and incubated overnight at 4°C with a 16-fold excess of prey proteins (total cell lysate). The next day, the beads were washed three times with TBS. Elution of bait and bound prey proteins was performed by adding 3xFLAG peptides according to the manufacturer’s instructions and incubating for 30 min at 4°C.

### TUBEs pull-down assays

THP-1-derived macrophages were treated with 1.6 µM rIpaH7.8 and respective controls together with 1µg/mL LPS (Invitrogen) and 10 µM MG132 (Sigma-Aldrich). After 6 hours of incubation, the cells were treated with 10 µM Nigericin (Invitrogen) to induce Gasdermin D cleavage and inflammasome formation for 30 min. The cells were washed with PBS, harvested by scraping the cells in 500 µL RIPA lysis buffer [25 mM Tris-HCl, pH 8.0; 137 mM NaCl; 0.1% (w/v) SDS; 0.5% (w/v) Na-deoxycholate; 10% (v/v) glycerol; 1% (v/v) NP-40; Complete protease inhibitor cocktail (Roche)]. The lysates were incubated on ice for 30 min and subsequently were clarified by centrifugation (20,000 x g, 30 min, 4°C), and equal amounts of protein were incubated with anti-ubiquitin-TUBE agarose beads (LifeSensors, Malvern, PA, USA) for 2 h at 4°C [14]. TUBE pull-downs were performed according to the manufacturer’s instructions. Briefly, the beads were washed three times with 1% Triton X-100 in TBS before the bound proteins were eluted using the provided elution buffer. The samples were divided into two equal portions; to one, 0.5 µg of USP2core enzyme was added, while the other was supplemented with the appropriate buffer. Both samples were incubated for 1 h at 37°C, and the reactions were terminated by the addition of 4x Laemmli buffer.

### ASC::GFP based speck formation analysis

Inflammasome speck formation analysis was adapted from Sester, 2015 [62]. Briefly, THP-1 cells expressing an ASC::GFP fusion protein were pre-stimulated with 1 µg/mL LPS to induce robust expression of the fusion construct and co-treated with rIpaH7.8 or rIpaH7.8ΔC to assess their effect on inflammasome formation. In addition, Z-VAD-FMK (50 µM; #S7023, Selleckchem), a PAN-caspase inhibitor, was applied to prevent the cells from pyroptotic cell death. Speck formation was triggered by the addition of 10 µM Nigericin (#tlrl nig, Invivogen) and analyzed by flow cytometry after 3 h. Speck-positive and speck-negative cells were distinguished based on their GFP signal profile in a pulse-width vs. pulse-height plot. A reduced width indicated a condensed GFP signal consistent with ASC speck formation.

### Analysis of IL-1β release

Cells were preincubated as indicated, either cells only (Medium), LPS (1µg/ml), or LPS with rIpaH7.8 (1.6 µM) or catalytically deficient IpaH7.8ΔC for 6 h. Inflammasome activation and Gasdermin D cleavage were triggered by the addition of 10 µM Nigericin (#tlrl-nig, Invivogen) for the indicated time. Cell-free supernatants were analyzed using human IL-1β ELISA MAX™ Deluxe Set (#437004, Biolegend) according to the manufacturer’s protocol. Samples were analysed in duplicates.

### RNA Sequencing and analysis

Primary human monocytes (1-day old) were stimulated with LPS (1 µg/mL) and incubated for six hours with recombinant IpaH7.8 or IpaH7.8ΔC (1.6 µM). Control samples consisted of cells treated only with medium or LPS. Total RNA was extracted using the Monarch® Total RNA Miniprep Kit (New England Biolabs). Further processing of the total RNA, including Poly-A enrichment, library preparation, and RNA sequencing, was performed using the Illumina NovaSeq X Plus platform with 150 bp paired-end reads (PE150) by Novogene (Munich, Germany). Bioinformatic analysis, including quality control, mapping, gene expression quantification, and differential gene expression analysis, was also performed by the sequencing provider. Genes were considered to be relevantly affected by catalytically active rIpaH7.8 treatment if they were identified as DEGs in LPS + rIpaH7.8 vs. LPS + rIpaH7.8ΔC as well as in at least one additional comparison (LPS + rIpaH7.8 vs. LPS or LPS + rIpaH7.8 vs. Untreated; Scheme see S4C Fig). These candidate genes were subjected to Gene Ontology (GO) enrichment analysis. Gene symbols were first mapped to Entrez Gene identifiers using the organism-specific annotation package org.Hs.eg.db, followed by enrichment analysis using the clusterProfiler package (v4.14.6) with the enrichGO function. Results were visualized as gene–concept networks (cnetplots) to illustrate the relationships between enriched GO terms and associated genes.

### Kinase activity profiling

The kinase activity profiling of 1-day-old primary human monocytes was performed using the PamChip assay, which quantifies kinase activity in cell and tissue lysates by measuring the phosphorylation of immobilized peptide substrates on PamChip® microarrays. Tyrosine and serine/threonine kinase activity profiling was conducted using a PamStation®12 (PamGene International B.V., ’s Hertogenbosch, Netherlands) with PamChip® peptide arrays according to the manufacturer’s instructions. Primary human monocytes were stimulated with LPS (1 µg/mL) and incubated for 6 h with recombinant rIpaH7.8 or rIpaH7.8ΔC (1.6 µM). Control samples consisted of cells treated only with medium or LPS. Following stimulation, the cells were lysed using M-PER™ Mammalian Protein Extraction Reagent (Thermo Fisher), supplemented with Halt™ Protease Inhibitor Cocktail (100x) and Halt™ Phosphatase Inhibitor Cocktail (100x) (Thermo Fisher). Lysates were centrifuged at 10,000 x g for 15 min at 4°C, and the cleared supernatants were aliquoted and stored at -80°C until further analysis. Protein concentrations were determined using Pierce™ BCA assay (Thermo Fisher Scientific) according to the manufacturer’s instructions. For kinase activity profiling, 5 µg of protein extract was used for the protein tyrosine kinase (PTK) protocol (v06), and 1 µg for the serine/threonine kinase (STK) protocol (v11) using Evolve software (PamGene). Signal intensities and correlation to kinase activity were analyzed using BioNavigator by Tercen.

To model kinase signaling networks, the UniProt IDs of parent proteins corresponding to the identified phosphosites were uploaded to MetaCore (Thomson Reuters), facilitating pathway analysis regarding regulatory kinase activity in primary human monocytes.

### Integrated Transcriptome and Kinome anaylsis

To identify cellular pathways, signaling hubs, and interaction networks associated with NEL-dependent effects across transcriptomic and kinome datasets, selected candidates were subjected to downstream enrichment analyses. For kinome data, kinases with altered inferred activity scores and phosphotargets exhibiting changes in relative phosphorylation signal intensities, as determined by PamGene PTK and STK arrays, were selected using criteria analogous to those applied for differentially expressed genes (S4C Fig): Candidates were considered as relevant (NEL-dependent effect) if they showed significant differences in rIpaH7.8 vs. rIpaH7.8ΔC upon LPS stimulation and additionally in at least one of the comparisons LPS stimulated and rIpaH7.8 treated vs. LPS alone or rIpaH7.8 vs. Untreated in inferred kinase activity or relative phosphorylation intensity. Resulting identifiers were, together with relevant gene-candidates subjected to KEGG-pathway enrichment analysis using the Enrichr web platform with the KEGG database to identify significantly enriched pathways [63]. To further explore functional relationships, protein-protein association networks were constructed using the STRING database (v11.5) [64]. Networks were generated using default parameters, incorporating known and predicted interactions derived from experimental evidence, curated databases, co-expression, and text mining. To facilitate the interpretation of identified pathways in the context of previously described IpaH7.8-associated processes, literature-curated factors were manually incorporated into the network.

### Measurement of intracellular reactive oxygen species (ROS)

Intracellular reactive oxygen species (ROS) production was quantified using the DCFDA/H₂DCFDA Cellular ROS Assay Kit (Abcam, Cambridge, UK; ab113851) according to the manufacturer’s instructions with minor modifications. Primary human monocytes were isolated from peripheral blood of healthy donors or thawed from cryopreserved stocks and seeded at a density of 2 × 10⁶ cells/mL in 6-well plates (3 mL per well). Cells were treated with lipopolysaccharide (LPS; 1 µg/mL) alone or in combination with recombinant rIpaH7.8 or rIpaH7.8ΔC (1.6 µM) overnight, as indicated. Medium-treated cells served as negative controls. Following stimulation with 150nM PMA for 2h, cells were collected, washed once with PBS, and incubated with DCFDA (20 µM) diluted in assay buffer for 30 min at 37 °C in the dark. After staining, cells were washed once with 1x buffer and resuspended in 1x supplemented buffer containing 10% heat-inactivated fetal calf serum (FCS). Cells were then transferred to black 96-well plates with clear bottoms (triplicates, 150 µL per well).

For positive controls, cells were treated with hydrogen peroxide (H₂O₂) immediately prior to measurement. Fluorescence was measured using a CLARIOstar microplate reader (BMG Labtech) pre-warmed to 37 °C at an excitation/emission of 485/535 nm, corresponding to oxidized DCF. Gain settings were adjusted using positive control wells to achieve ∼80% of maximal signal. Blank wells containing buffer only were used for background subtraction. ROS levels are presented as mean fluorescence relative fluoresecne units (RFU) after background correction.

### Structural Modeling and Surface Charge Visualization

Protein structure predictions were performed using the AlphaFold Server (Google DeepMind) [65]. The resulting PDB files were visualized in UCSF Chimera (Chimera v1.1.9 (Resource for Biocomputing, Visualization, and Informatics, UCSF with support from NIH P41-GM103311) [66], and electrostatic surface potentials were rendered using the Coulombic surface coloring tool, accessed through Tools → Surface/Binding Analysis. Theoretical net charges of the selected peptide and protein segments were calculated at pH 5.5, 6.5, and 7.4 using the ProtPi Peptide Tool (www.protpi.ch/Calculator/PeptideTool v2.2.29.152). Calculations were based on unmodified primary sequences and standard pKa values for ionizable residues.

### Ethics Statement

Human primary monocytes were isolated from leukocyte reduction system (LRS) chambers obtained from fresh platelet apheresis donations of anonymized healthy adult volunteers. Blood donations were collected at the University Hospital Münster in accordance with institutional guidelines and national regulations. Written informed consent was obtained from all donors prior to donation. Due to the use of fully anonymized donor material, no specific ethical approval was required.

## Acknowledgments

This study was supported by grants from the Deutsche Forschungsgemeinschaft (DFG, SFB1009 TP B03 to Christian Rüter, DFG, KFO342 P10 and SFB DECIDE B03 to Petra Dersch. The funders had no role in study design, data collection, and interpretation, or the decision to submit the work for publication. We thank Judith Austermann (Institute of Immunology, University of Münster) for expert training and support in the isolation of primary human monocytes. We would like to thank our colleagues at the Institute of Infectiology - ZMBE for fruitful discussions and their valuable contributions. This work is part of the doctoral thesis of T.K. Author contributions: TK, INY, and CR designed the study; TK, INY, SN, BK, YB, JF, and YT performed experiments; CR and PD supervised the study and raised funding; TK and CR wrote the manuscript. We thank Ellie Schmidt for thorough proofreading.

## Supplemental Figures

**Supplemental Figure S1.**
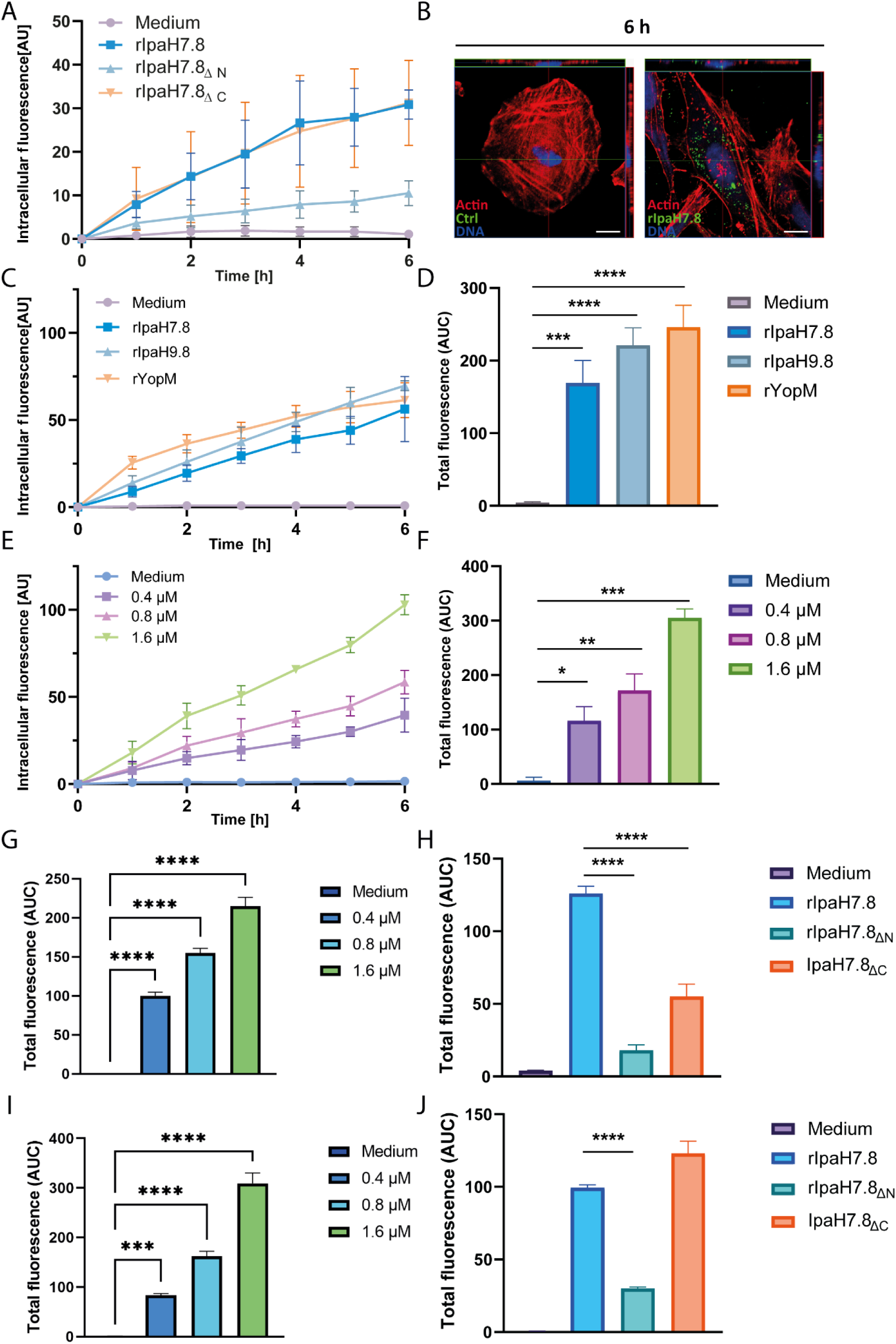
T3SS-independent uptake of rIpaH7.8 in epithelial and immune cell lines. Flow cytometry-based quantification of intracellular fluorescence in HeLa (A, C-F), HEK293 (G, H) and THP-1 macrophages (I, J) incubated with increasing concentrations (0.4-1.6 µM) of FITC-labeled rIpaH7.8 (E, F, G, I) and rIpaH7.8/rYopM (C, D) or with 0.4 µM FITC-labeled rIpaH7.8ΔN or rIpaH7.8ΔC for up to 6 h (A, C, H, J). Medium-only samples served as controls. Data are shown as mean ± SD of three independent experiments; AU, arbitrary units. Area under the curve (AUC) values are shown for visualization. Statistical analysis by one-way ANOVA with Tukey’s post hoc test; ns: not significant, *p < 0.05, **p < 0.01, ***p < 0.001, ****p < 0.0001 vs. medium. (B) Representative fluorescence microscopy images of HeLa cells incubated with 0.8 µM FITC-labeled rIpaH7.8 for 6 h. Actin (red), FITC-rIpaH7.8 (green), nuclei (blue). Scale bar, 10 µm. Images show merged channels from a single optical section.

**Supplemental Figure S2.**
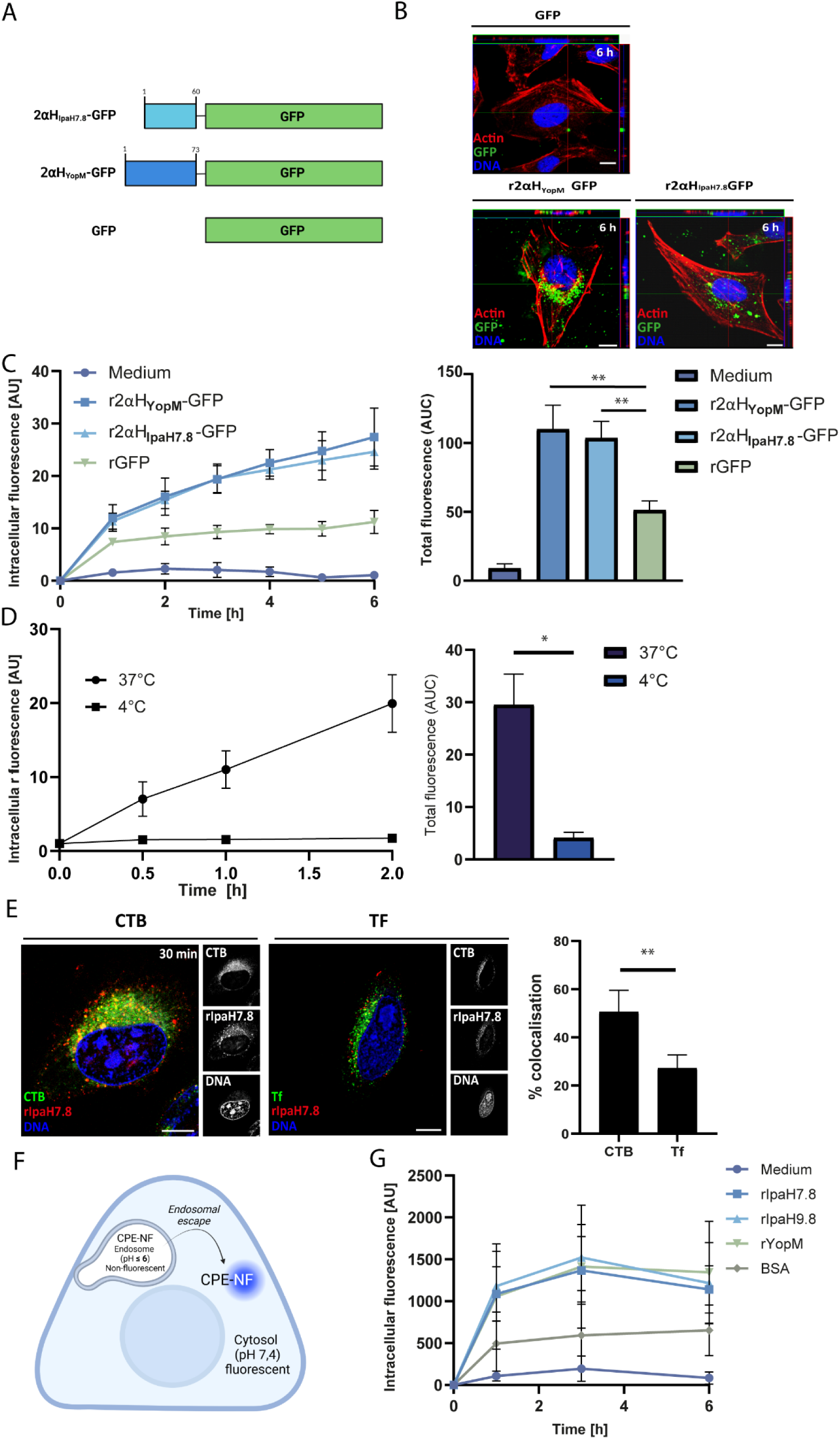
Uptake and cytosolic delivery of IpaH7.8 GFP fusion constructs. (A) Schematic overview of GFP fusion constructs: 2αHIpaH7.8-GFP and 2αHYopM-GFP, containing the N-terminal domains of IpaH7.8 or YopM; rGFP served as control. (B) Representative fluorescence microscopy images of HeLa cells incubated with 0.8 µM FITC-labeled rGFP, 2αHYopM-GFP, or r2αHIpaH7.8-GFP for 6 h. Actin (red), protein (green), nuclei (blue). Scale bar, 10 µm. (C) Flow cytometry quantification of intracellular GFP fluorescence after incubation with 0.4 µM FITC-labeled proteins. Data show mean ± SD (n = 3); AUC values are indicated. One-way ANOVA with Tukey’s post hoc test (ns, *p < 0.05 vs. rGFP). (D) Temperature-dependent uptake of Alexa Fluor 488-labeled rIpaH7.8 at 37 °C or 4 °C for up to 3 h (AUC; n = 3; paired t-test, *p < 0.01). Co-localization of Cy3-rIpaH7.8 with Cholera Toxin B-subunit CTB-FITC (lipid rafts) or transferrin-Alexa Fluor 488 (Tf) after 30 min. Quantification from six images are shown (paired t-test, **p < 0.01). Scale bar, 10 µm. (F) Experimental scheme for detecting cytosolic localization using the pH-sensitive dye 5(6)-carboxynaphthofluorescein (NF). (G) Flow cytometry of HeLa cells incubated with 0.4 µM NF-labeled rIpaH7.8, rIpaH9.8, rYopM, or BSA for up to 6 h. Data show mean ± SD (n = 3); AU, arbitrary units.

**Supplemental Figure S3.**
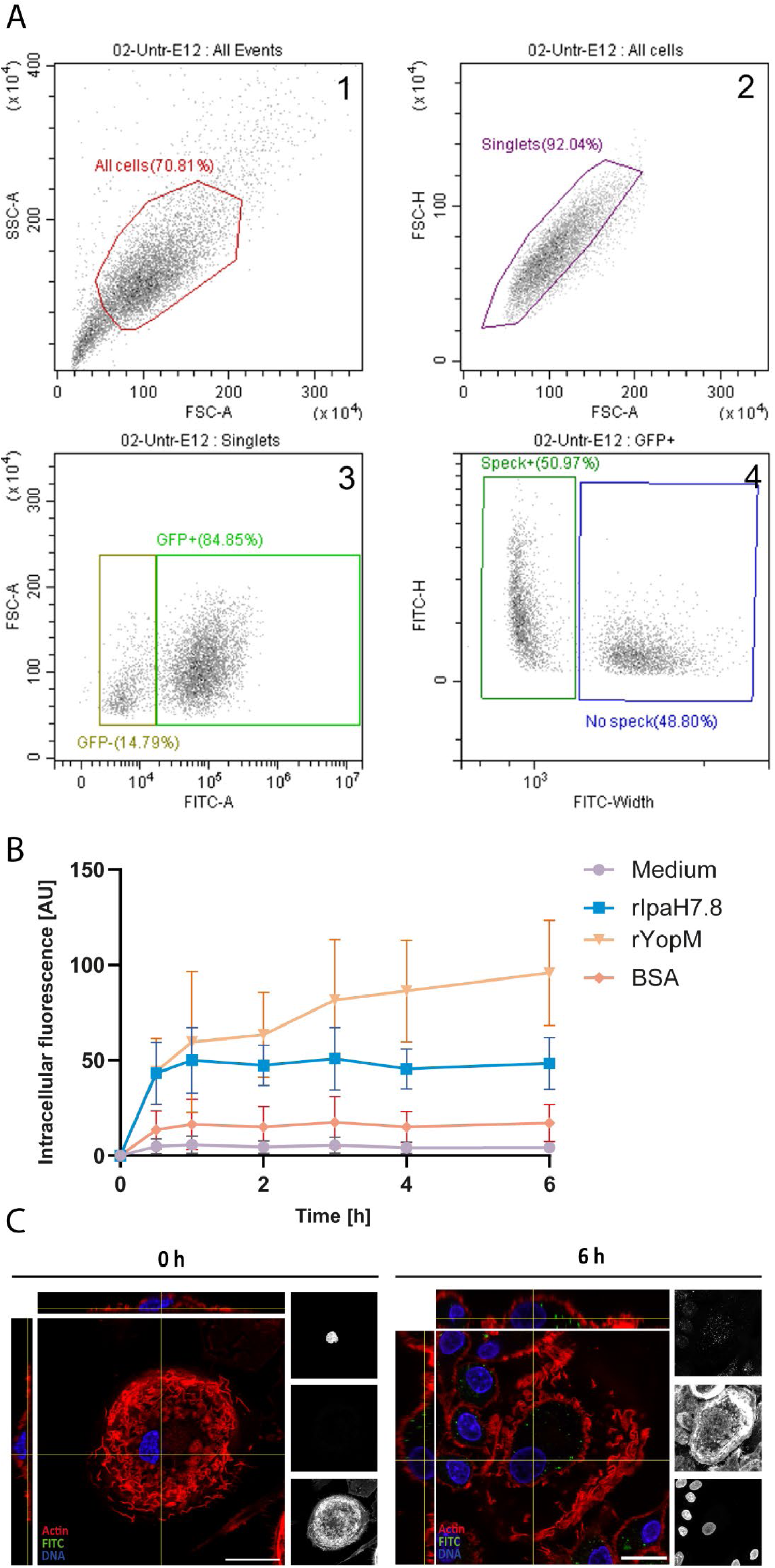
ASC speck gating strategy and uptake of rIpaH7.8 in primary human monocytes. (A) Flow cytometry gating strategy for ASC speck formation in THP-1 cells expressing ASC::GFP. Cells were gated for singlets and GFP-positive events, followed by discrimination of speck-positive and speck-negative populations based on FITC height versus width. (B) Uptake of FITC-labeled rIpaH7.8, rYopM, or BSA by primary human monocytes quantified by flow cytometry after incubation for up to 6 h. Medium-only served as a negative control. n = 7 donors; mean ± SD; AU, arbitrary units. (C) Confocal microscopy of primary human monocyte-derived macrophages incubated with FITC-labeled rIpaH7.8 (1.6 µM, 6 h). Actin (red), rIpaH7.8 (green), nuclei (blue). Medium-only (0 h) served as control. Scale bar, 20 µm. Orthogonal and maximum projections show single optical sections (1 airy unit).

**Supplementary Figure S4.**
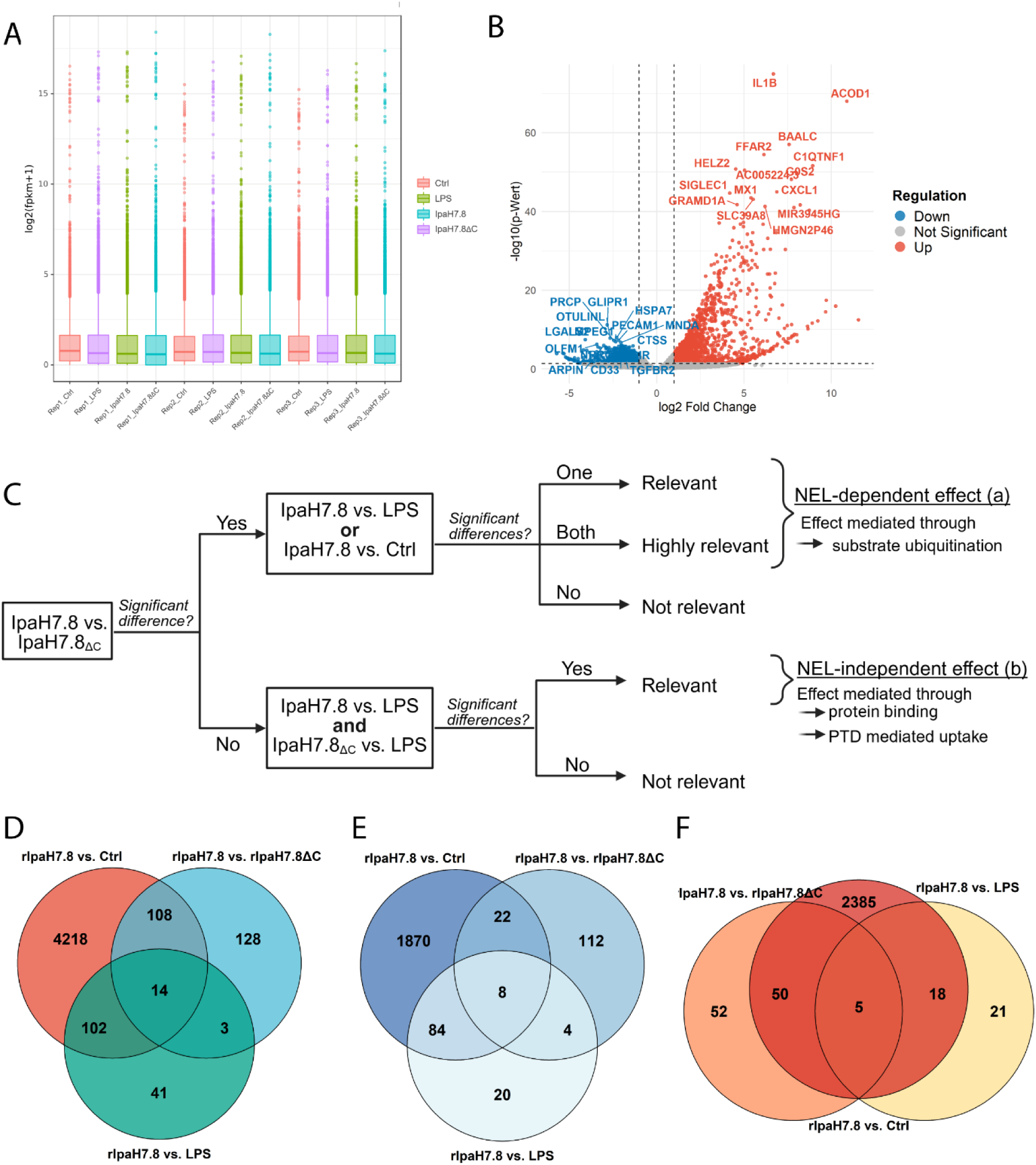
Transcriptomic quality control, differential expression, and analysis strategy. (A) Gene expression distributions across biological replicates for each condition (Ctrl, LPS, LPS + rIpaH7.8, LPS + rIpaH7.8ΔC), demonstrating comparable global expression profiles. (B) Volcano plot showing differential gene expression in LPS-treated versus untreated monocytes. (C) Schematic overview of the pairwise comparison strategy used to define NEL-dependent and NEL-independent gene regulation. NEL-dependent genes were defined as differentially expressed exclusively in cells treated with catalytically active rIpaH7.8, whereas NEL-independent genes showed comparable regulation in both rIpaH7.8- and rIpaH7.8ΔC-treated cells (p < 0.05). For clarity, LPS co-treatment present in all conditions is not depicted. (D-F) Venn diagram analyses of differentially expressed genes across the indicated comparisons (Fig 4C-E), showing total (D), downregulated (E), and upregulated (F) DEGs.

**Supplementary Figure S5.**
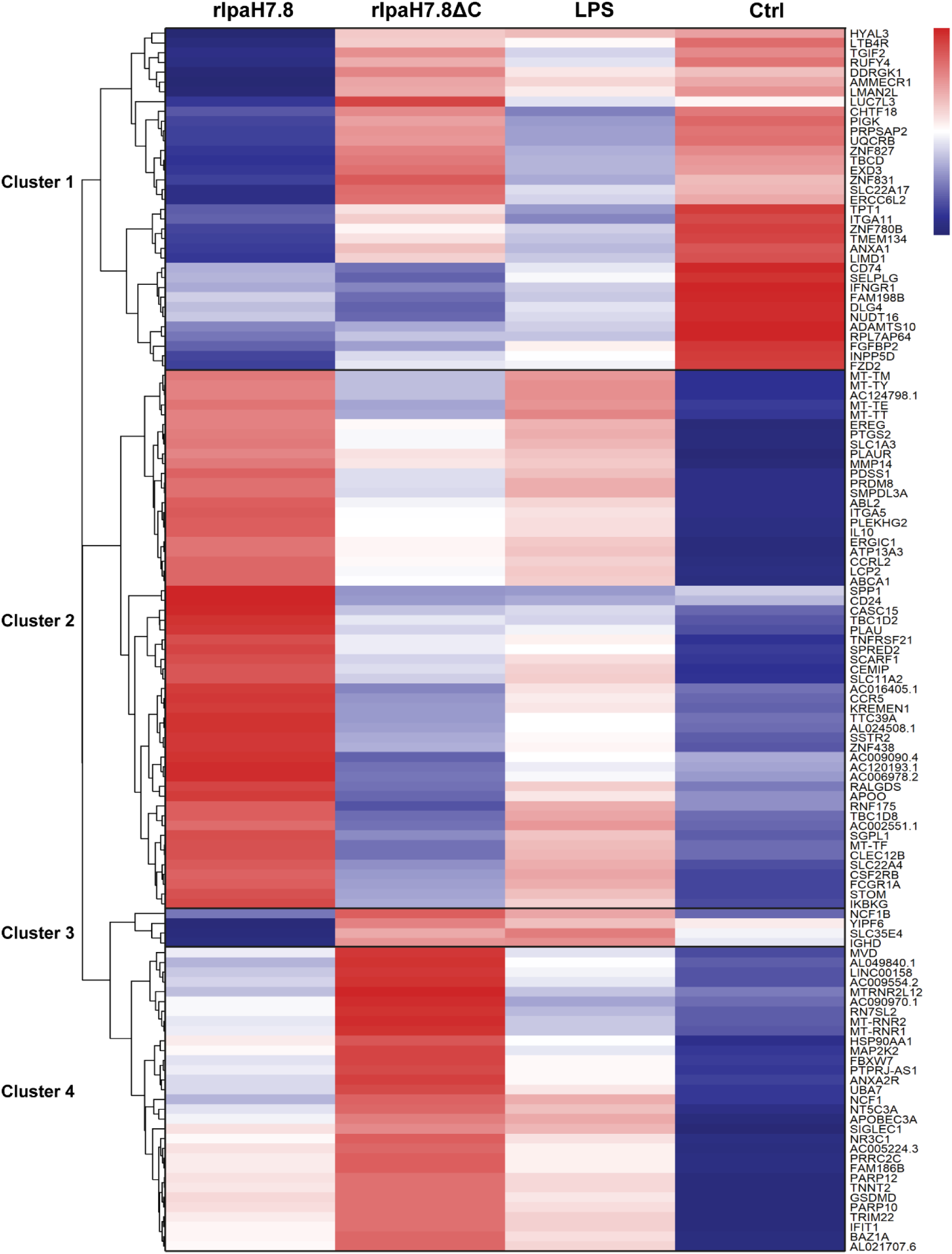
rIpaH7.8-dependent transcriptional signatures in LPS-stimulated monocytes. Heatmap of differentially expressed genes (DEGs) identified by the intersectional filtering strategy shown in Fig. S4D and summarized in Fig. 4F. Expression values are shown as Z-score-normalized FPKM values per gene to emphasize relative differences across conditions. Rows represent individual genes clustered by Euclidean distance, and columns correspond to the indicated treatment conditions (LPS + rIpaH7.8, LPS + rIpaH7.8ΔC, LPS only, and untreated control). Red indicates higher relative expression and blue indicates lower relative expression. Unsupervised clustering indicates four major gene expression patterns.

**Supplemental Figure S6.**
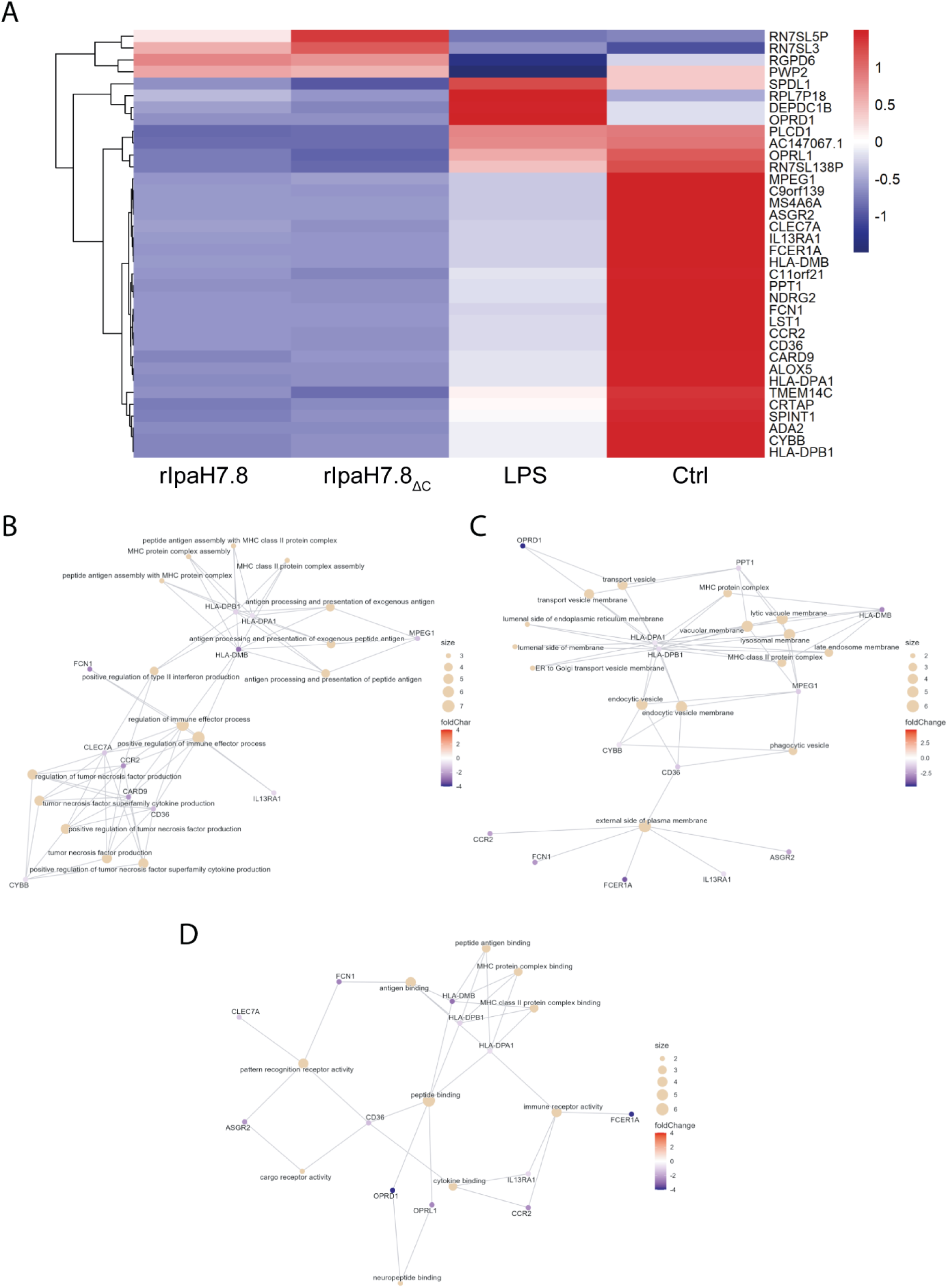
Transcriptomic changes independent of NEL activity. (A) Heatmap of NEL-independent differentially expressed genes identified by the intersectional strategy shown in Fig. S4C. Z-score-normalized FPKM values are shown for each gene (rows), clustered by Euclidean distance, across the indicated conditions (rIpaH7.8, rIpaH7.8ΔC, LPS, and untreated). Red indicates higher and blue lower relative expression. (B-D) GO enrichment analysis of NEL-independent overlapping DEGs displayed as cnet plots for (B) Biological Processes, (C) Cellular Components, and (D) Molecular Functions. GO terms are shown as central nodes, with connected gene nodes indicating membership.

**Supplemental Figure S7.**
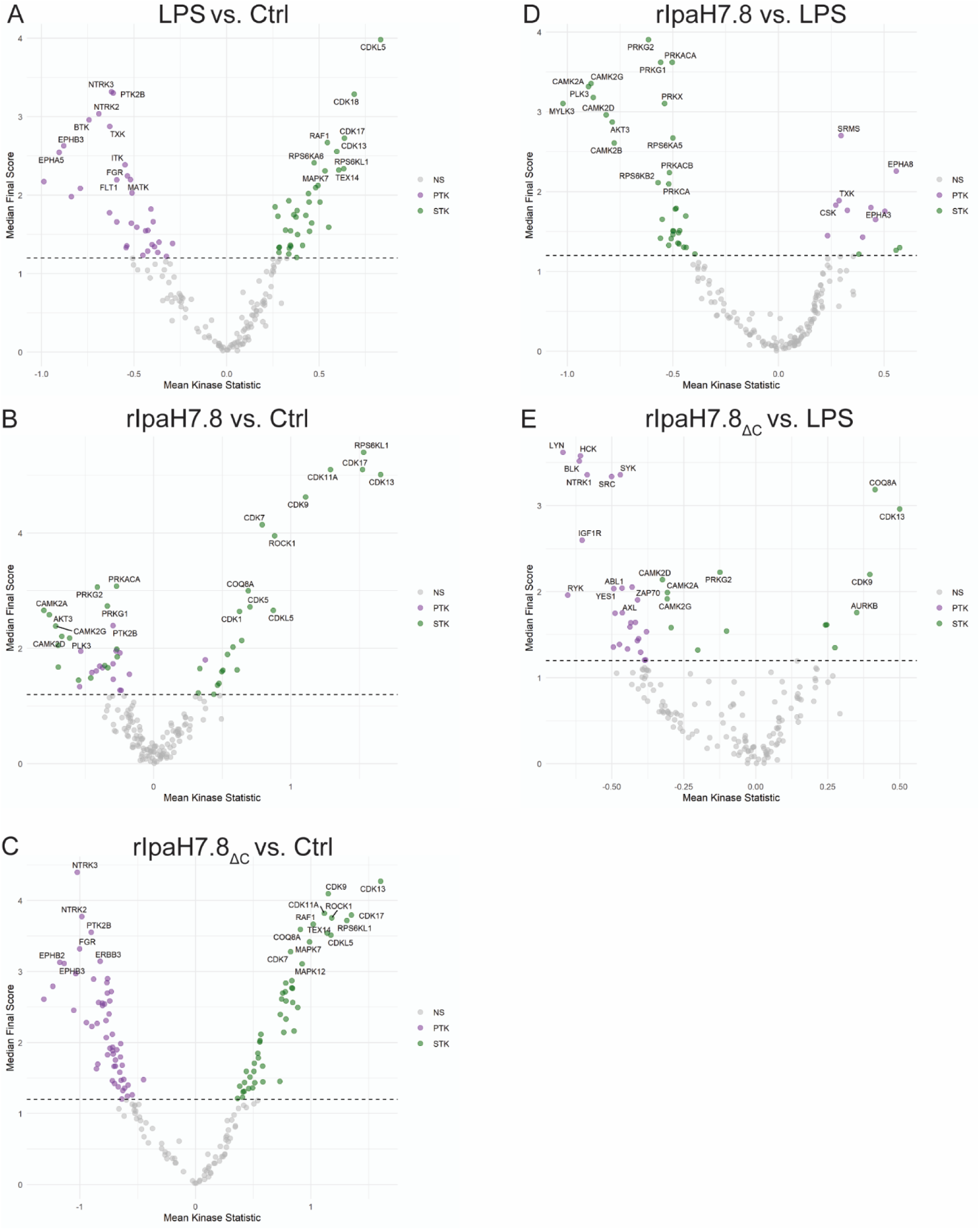
Kinome profiling reveals rIpaH7.8-dependent modulation of LPS-induced kinase signaling. Primary human monocytes were analyzed using PamGene kinase arrays to infer upstream kinase activity changes under the indicated conditions. (A) Volcano plot showing inferred kinase activity changes following LPS stimulation compared to untreated control cells, revealing broad alterations in PTK and STK activities. (B) Kinase activity changes in monocytes treated with catalytically active rIpaH7.8 compared to control cells, indicating direct effector-mediated modulation of selected kinase families. (C) Kinase activity profile of cells treated with the catalytically inactive rIpaH7.8ΔC mutant compared to control, demonstrating partially overlapping but distinct, NEL-independent effects. (D) Comparison of LPS-stimulated cells treated with rIpaH7.8 versus LPS alone, showing partial reversal of LPS-induced kinase attenuation and restoration of specific immune-related kinase activities. (E) Comparison of LPS-stimulated cells treated with rIpaH7.8ΔC versus LPS alone, highlighting the requirement of the NEL domain for effective modulation of LPS-driven kinase responses. Each dot represents a kinase-associated peptide; significantly regulated peptides are highlighted (PTK, purple; STK, green), while non-significant peptides are shown in grey. The dashed line indicates the significance threshold (median final score > 1.2).

**Supplemental Figure S8.**
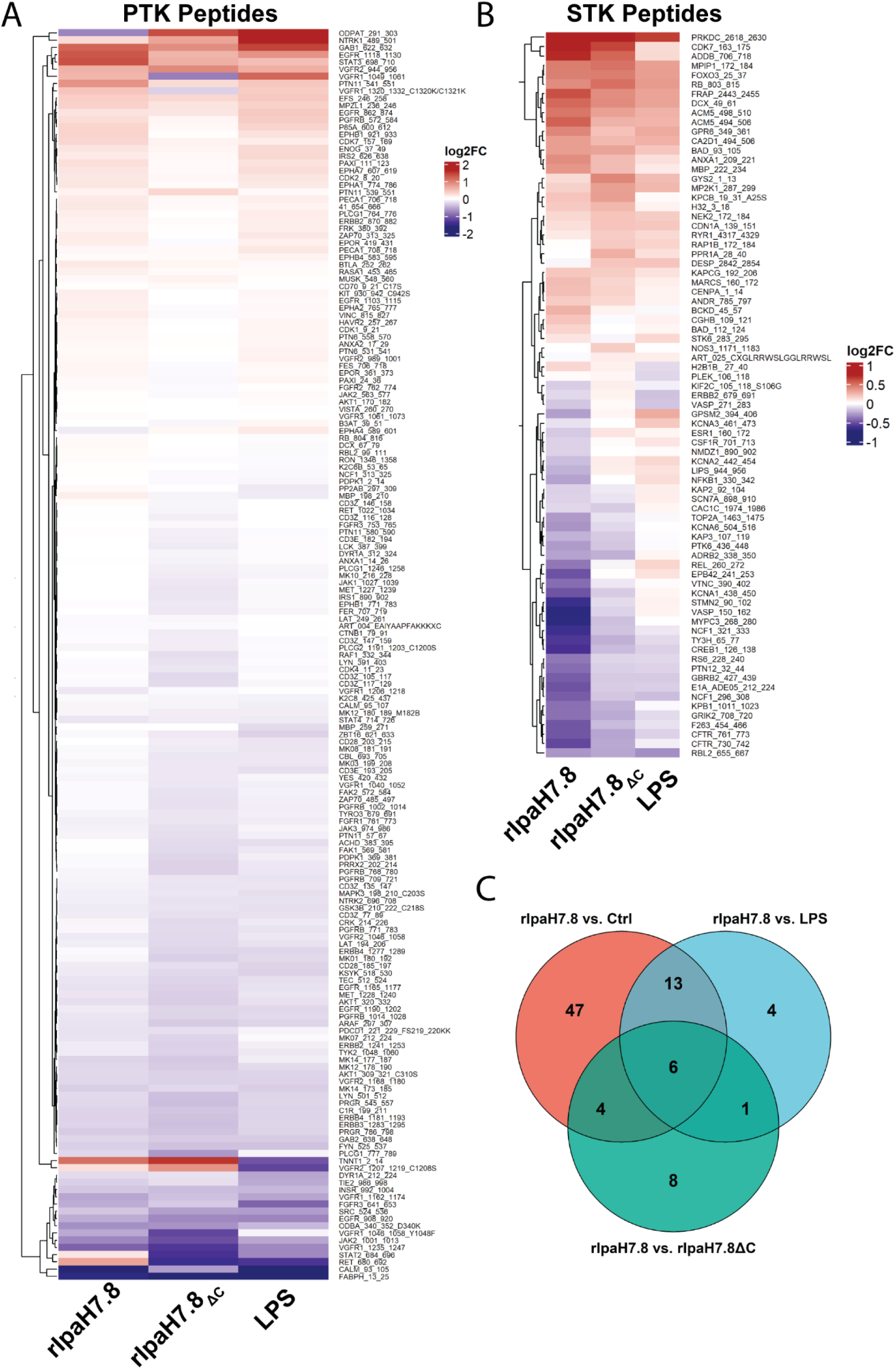
Relative peptide phosphorylation profiles in response to rIpaH7.8. Primary human monocytes were stimulated with LPS (1 µg/mL) alone or in combination with rIpaH7.8 or rIpaH7.8ΔC (1.6 µM) for 6 h. Cell lysates were analyzed using PamGene kinase microarrays. Clustered heatmaps display log2 fold changes (log2FC) in phosphorylation of peptide substrates derived from protein tyrosine kinases (PTK, A) or serine/threonine kinases (STK, B) relative to untreated controls. Blue indicates reduced and red increased phosphorylation. Data represent three biological replicates, each consisting of pooled samples from three independent donors. (C) Venn diagram analyses of differentially phosphorylated peptides across the indicated comparisons.

**Supplemental Figure S9.**
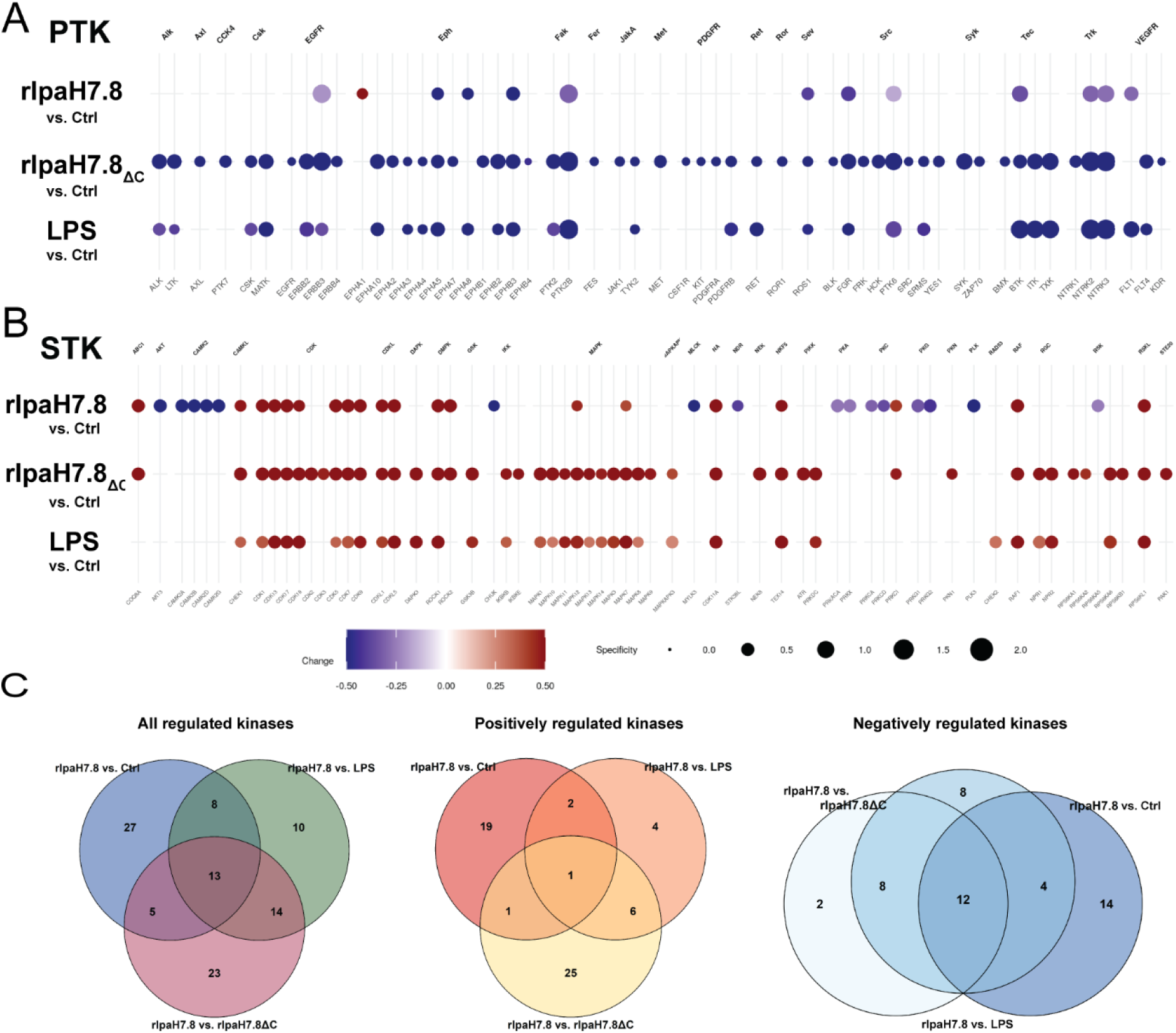
Kinome profiling of rIpaH7.8-treated human monocytes. (A) PTK and STK (B) activity across conditions. Color indicates relative activity changes (blue, decreased; white, unchanged; red, increased); node size reflects substrate specificity. (C) Venn diagrams depicting kinases significantly affected by rIpaH7.8 across comparisons, including all, positively regulated, and negatively regulated kinases.

**Supplemental Figure S10.**
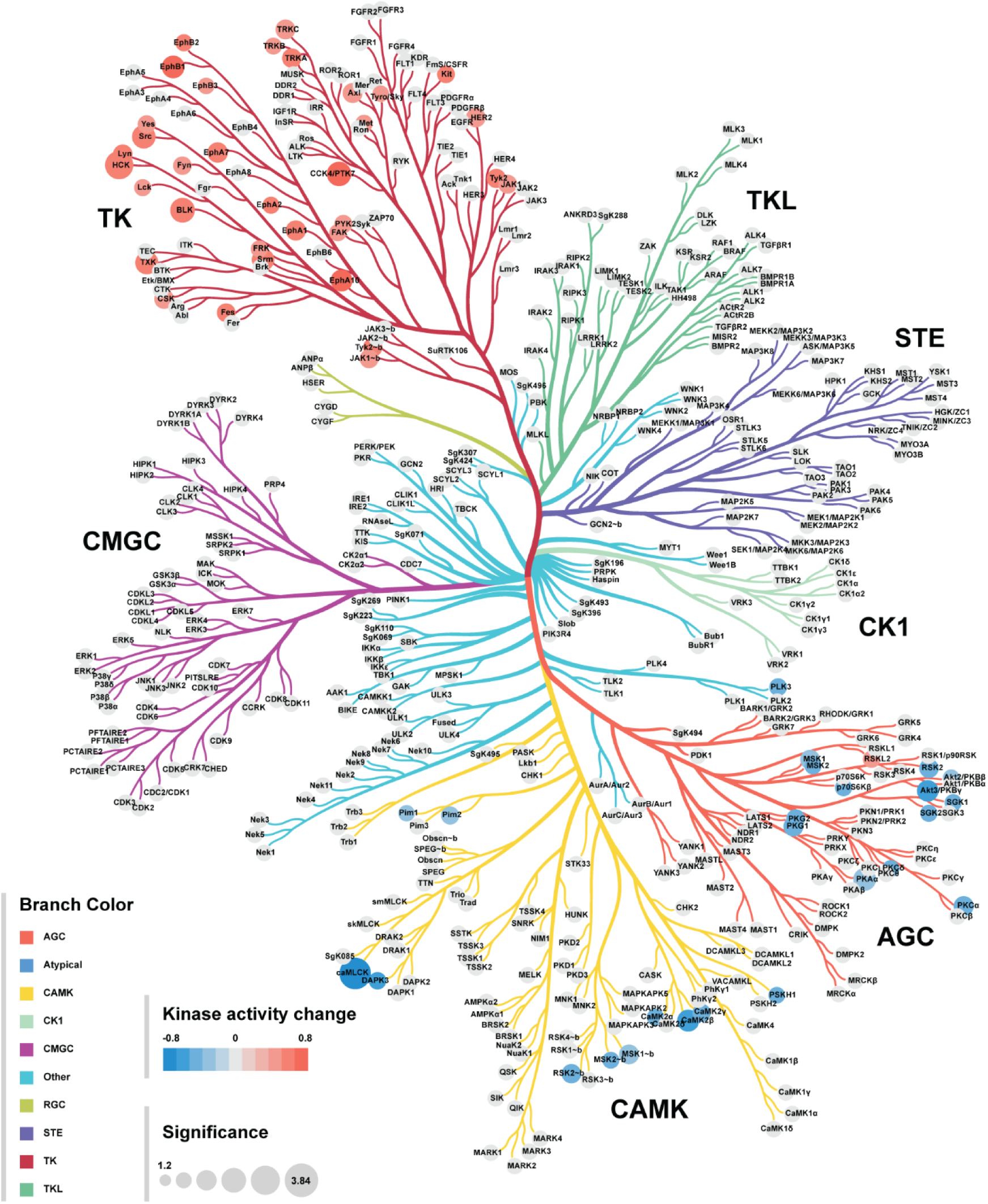
Kinome tree visualization of rIpaH7.8-induced kinase modulation. Differentially active kinases identified by upstream kinase analysis were mapped onto the human kinome tree. Node size reflects predicted kinase activity scores, and node color indicates the direction of regulation in rIpaH7.8-treated cells relative to rIpaH7.8ΔC. Affected kinases cluster across multiple families, including AGC, CMGC, and TKL, indicating broad catalytic activity-dependent reprogramming of host kinase signaling.

**Supplemental Figure 11:**
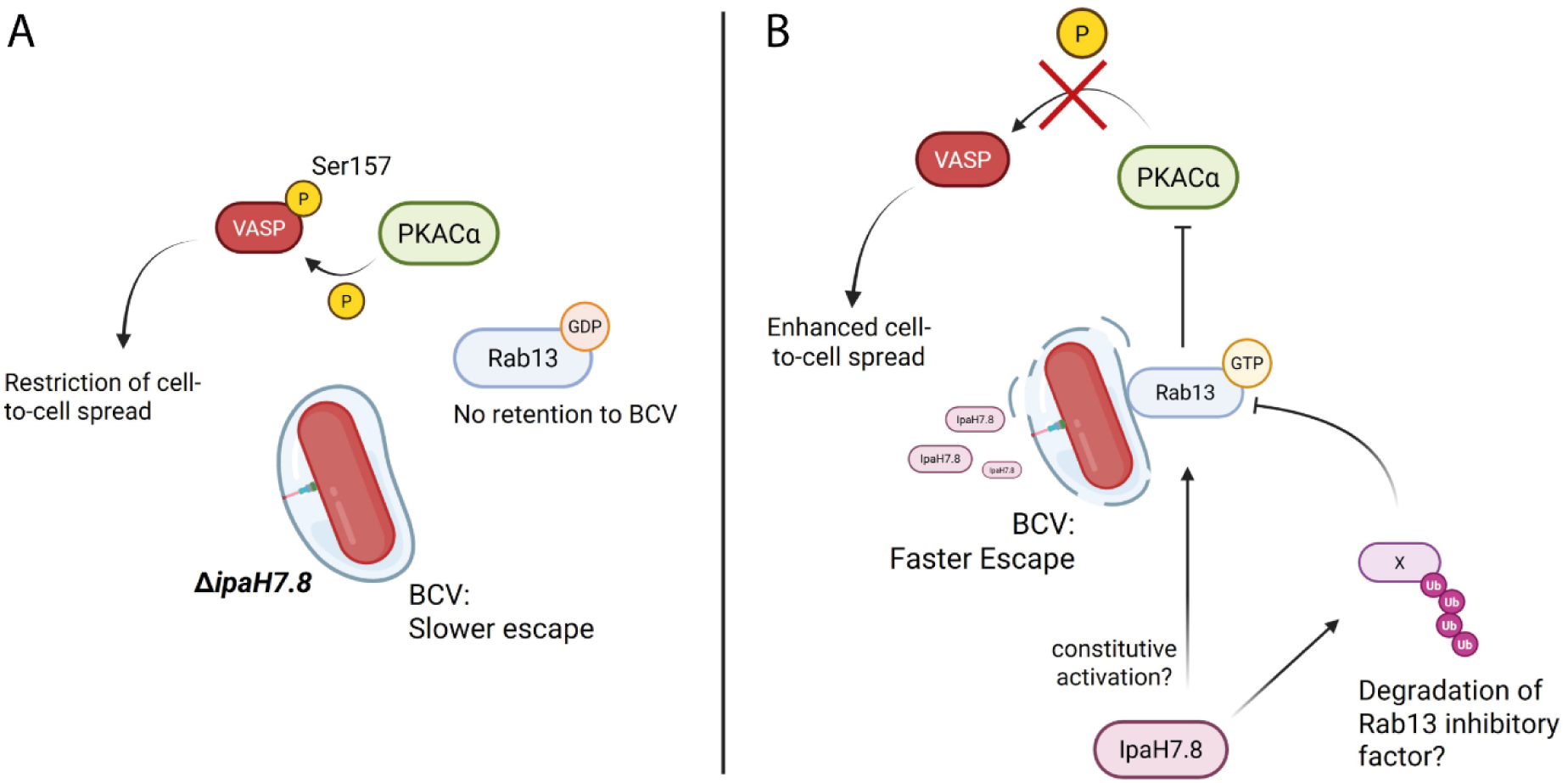
Proposed mechanism of IpaH7.8 mediated promotion of vacuolar escape of *Shigella*. (A) In absence of IpaH7.8 mediated Rab13-retention, VASP Ser157 phosphorylation is conducted by PKACα, which in turn restricts cell-to-cell dissemination. (B) In presence of IpaH7.8, Rab13 is kept in its, active GTP bound form, leading to inhibition of PKACα and thus prevents VASP phosphorylation. This abolishes cell-to-cell spread restriction and in a so far unclear mechanism promotes vacuolar disassembly.

**Supplemental Table 1:** Numeric values of most regulated genes by rIpaH7.8 compared to LPS stimulation alone.

| Gene_ID | Name | IpaH7.8 vs Ctrl |  | IpaH7.8 vs LPS |  | IpaH7.8 vs IpaH7.8<sub>ΔC</sub> |  | LPS vs. Ctrl |  |
| --- | --- | --- | --- | --- | --- | --- | --- | --- | --- |
|  |  | log2FC | p-value | log2FC | p-value | log2FC | p-value | log2FC | p-value |
| ENSG00000272398 | CD24 | 2.347 | 0.019 | 2.565 | 0.021 | 2.958 | 0.022 | -0.180 | 0.887 |
| ENSG00000118785 | SPP1 | 1.535 | 0.020 | 1.966 | 5.88E-03 | 1.997 | 0.022 | -0.389 | 0.537 |
| ENSG00000272168 | CASC15 | 3.660 | 7.96E-07 | 1.595 | 0.013 | 1.573 | 0.042 | 2.072 | 0.031 |
| ENSG00000198369 | SPRED2 | 2.896 | 4.89E-11 | 0.853 | 0.043 | 0.920 | 0.038 | 2.046 | 9.89E-05 |
| ENSG00000095383 | TBC1D2 | 1.132 | 5.52E-04 | 0.654 | 0.038 | 0.604 | 0.047 | 0.480 | 0.234 |
| ENSG00000043462 | LCP2 | 1.983 | 7.42E-16 | 0.330 | 0.218 | 0.577 | 0.010 | 1.661 | 5.96E-07 |
| ENSG00000161638 | ITGA5 | 1.487 | 1.08E-09 | 0.401 | 0.144 | 0.500 | 0.041 | 1.092 | 3.96E-04 |
| ENSG00000135046 | ANXA1 | -0.987 | 0.003 | -0.408 | 0.268 | -0.773 | 0.033 | -0.591 | 0.030 |
| ENSG00000168918 | INPP5D | -1.619 | 1.17E-11 | -0.969 | 0.003 | -0.817 | 8.88E-03 | -0.661 | 0.032 |
| ENSG00000158517 | NCF1 | 0.723 | 0.001 | -0.580 | 0.020 | -0.819 | 8.31E-05 | 1.309 | 3.66E-06 |
| ENSG00000182487 | NCF1B | 0.131 | 0.721 | -0.769 | 0.041 | -0.977 | 0.010 | 0.900 | 0.004 |
| ENSG00000211898 | IGHD | -0.861 | 0.075 | -1.333 | 8.70E-03 | -1.304 | 0.038 | 0.477 | 0.253 |
| ENSG00000181704 | YIPF6 | -1.197 | 0.038 | -1.388 | 2.29E-02 | -1.582 | 0.020 | 0.189 | 0.678 |
| ENSG00000114988 | LMAN2L | -1.781 | 5.58E-03 | -1.520 | 0.040 | -1.764 | 0.017 | -0.276 | 0.599 |
| ENSG00000188282 | RUFY4 | -2.003 | 1.46E-04 | -1.412 | 0.035 | -1.905 | 0.008 | -0.606 | 0.219 |
| ENSG00000180340 | FZD2 | -3.031 | 2.30E-06 | -2.063 | 0.013 | -1.944 | 0.032 | -0.981 | 0.056 |
| ENSG00000128000 | ZNF780B | -3.144 | 1.07E-06 | -1.872 | 0.038 | -2.246 | 0.014 | -1.273 | 0.017 |
| ENSG00000100036 | SLC35E4 | -2.320 | 0.083 | -3.006 | 0.021 | -2.838 | 0.045 | 0.689 | 0.379 |
| ENSG00000137809 | ITGA11 | -4.001 | 2.56E-03 | -1.257 | 0.562 | -3.282 | 0.040 | -2.811 | 7.43E-03 |
| ENSG00000186792 | HYAL3 | -4.783 | 4.21E-03 | -4.666 | 0.015 | -4.604 | 0.022 | -0.180 | 0.851 |
| ENSG00000213876 | RPL7AP64 | -7.319 | 1.79E-08 | -5.037 | 5.76E-03 | -4.925 | 0.023 | -2.366 | 7.76E-04 |

**Supplemental Table S2: Contrasts of peptide phosphorylation analysis**

**(attached as supplemental file)**

**Supplemental Table 3:** List of relevant kinases significantly regulated by rIpaH7.8 treatment (NEL dependent; IpaH7.8 vs. IpaH7.8 _ΔC_).

| Kinase | Uniprot-ID | Median Final score<br>(Specificity) | Mean Kinase Statistic<br>(Kinase activity change) | Chip |
| --- | --- | --- | --- | --- |
| MYLK3 | Q32MK0 | 3.854 | -0.874 | STK |
| PTK7 | Q13308 | 2.601 | 0.719 | PTK |
| TXK | P42681 | 2.387 | 0.546 | PTK |
| PRKACA | P17612 | 2.322 | -0.356 | STK |
| SRMS | Q9H3Y6 | 2.255 | 0.532 | PTK |
| CAMK2B | Q13554 | 2.179 | -0.582 | STK |
| EPHA10 | Q5JZY3 | 2.158 | 0.727 | PTK |
| AKT3 | Q9Y243 | 2.088 | -0.559 | STK |
| EPHA1 | P21709 | 1.863 | 0.598 | PTK |
| PRKCA | P17252 | 1.834 | -0.412 | STK |
| PRKG1 | Q13976 | 1.834 | -0.368 | STK |
| RPS6KA5 | O75582 | 1.793 | -0.344 | STK |
| CSK | P41240 | 1.785 | 0.488 | PTK |
| RPS6KA3 | P51812 | 1.724 | -0.411 | STK |
| PRKG2 | Q13237 | 1.631 | -0.367 | STK |
| FES | P07332 | 1.608 | 0.539 | PTK |
| CAMK2G | Q13555 | 1.525 | -0.453 | STK |
| PIM2 | Q9P1W9 | 1.512 | -0.342 | STK |
| PLK3 | Q9H4B4 | 1.427 | -0.423 | STK |
| RPS6KL1 | Q9Y6S9 | 1.379 | 0.488 | STK |
| CAMK2A | Q9UQM7 | 1.340 | -0.465 | STK |
| RPS6KB2 | Q9UBS0 | 1.333 | -0.378 | STK |
| SGK2 | Q9HBY8 | 1.332 | -0.415 | STK |
| SGK1 | O00141 | 1.324 | -0.405 | STK |
| PRKCD | Q05655 | 1.299 | -0.372 | STK |
| RPS6KA4 | O75676 | 1.285 | -0.437 | STK |
| PSKH1 | P11801 | 1.239 | -0.478 | STK |
| AKT2 | P31751 | 1.231 | -0.353 | STK |

**Supplemental Table 4:** List of relevant kinases significantly regulated by rIpaH7.8 treatment (NEL independent; IpaH7.8 vs. LPS). Full list of peptide regulation can be accessed at xxxDOI; Suppl_Data_S4 Tab

| Kinase | Uniprot-ID | Median Final score<br>(Specificity) | Mean Kinase Statistic<br>(Kinase activity change) | Chip |
| --- | --- | --- | --- | --- |
| PRKX | P51817 | 3.106 | -0.269 | STK |
| CAMK2D | Q13557 | 2.963 | -0.671 | STK |
| CDK13 | Q14004 | 1.265 | 1.655 | STK |

